# The ecology of cancer prevalence across species: Cancer prevalence is highest in desert species and high trophic levels

**DOI:** 10.1101/2022.08.23.504890

**Authors:** Stefania E. Kapsetaki, Zachary Compton, Shawn M. Rupp, Michael M. Garner, Elizabeth G. Duke, Amy M. Boddy, Tara M. Harrison, Athena Aktipis, Carlo C. Maley

**Affiliations:** Arizona Cancer Evolution Center, Arizona State University, Tempe, AZ, USA; Center for Biocomputing, Security and Society, Biodesign Institute, Arizona State University, Tempe, AZ, USA; School of Life Sciences, Arizona State University, Tempe, AZ, USA; Northwest ZooPath, Monroe, WA 98272, USA; Department of Clinical Sciences, North Carolina State University, Raleigh, NC, 27607 USA; Department of Anthropology, University of California Santa Barbara, CA, USA; Department of Psychology, Arizona State University, Tempe, AZ, USA; Exotic Species Cancer Research Alliance, North Carolina State University, Raleigh, NC, 27607 USA

**Author notes:** co-senior authors (Athena Aktipis, Carlo C. Maley).

**Keywords:** biome, cancer prevalence, comparative oncology, habitat, metabolism, trophic level

## Abstract

The ecology in which species live and evolve likely affects their health and vulnerability to diseases including cancer. Using 14,267 necropsy records across 244 vertebrate species, we tested if animals in low productivity habitats, with large habitat range, high body temperature and weight-inferred estimates of metabolic rates, and in high trophic levels (from lowest to highest: herbivores, invertivores, primary carnivores, and secondary carnivores) are linked with having increased prevalence of neoplasia. This study found that: (1) habitat productivity negatively correlated with the prevalence of malignancy and neoplasia across tissues, and malignancy and neoplasia in gastrointestinal tissues; (2) inferred metabolic rates negatively correlated with the prevalence of neoplasia; and (3) trophic levels positively correlated with malignancy and neoplasia prevalence in both mammals and non-mammals. However, only the correlations with trophic levels remained significant after Bonferroni corrections for multiple testing. There are several mechanisms that might explain these findings, including the biomagnification of carcinogens in higher trophic levels, as well as tradeoffs between cancer suppression versus reproduction and survival in low productivity environments.

## Introduction

Very little is currently understood about why some organisms evolve to be more susceptible to cancer than others, and what tradeoffs constrain the evolution of cancer suppression. Previous work has looked at the association between cancer prevalence and intrinsic organismal characteristics such as body size, longevity, placental invasiveness, and litter size^1–3^. Because the ecology of a species determines the selective pressures on that species, we hypothesised that aspects of a species ecology may force tradeoffs between cancer suppression and adaptations to the environment. This study investigates if there are associations between cancer prevalence and habitat productivity, habitat range, temperature- and weight-inferred estimates of metabolic rates, and trophic levels. For trophic levels, we classified species by their primary diet as herbivores (plant eaters), invertivores (invertebrate eaters), primary carnivores (herbivore and invertivore eaters), and secondary carnivores (eaters of primary carnivores).

The selective pressures in the ecology of a species shape all aspects of that species’ life history, including its susceptibility to cancer. Life history theory predicts that larger and longer-lived animals may have invested more energy in cancer defence mechanisms, e.g. more protection from mutations, and less energy in reproduction than smaller and shorter-lived animals^4^, and thus have lower cancer prevalence. Cancer researchers, however, predict that larger, longer-lived organisms should have more cancer, because they have more cells that could generate cancer over longer lifespans^5–8^. The observation that larger, longer-lived species do not seem to get more cancer is called Peto’s Paradox^1, 2, 9^. However, there is a life history trait, litter size, that is positively correlated with cancer prevalence across 29 mammals, indicating a possible tradeoff between offspring quantity and quality of somatic maintenance^1^.

### Cancer vulnerability should be associated with ecological variables

Low productivity habitats may select for species that are more susceptible to cancer through a variety of mechanisms including tradeoffs between investment in somatic maintenance versus reproduction and survival in resource limited environments, selection for lower metabolisms and meat-eating. Habitat productivity can be quantified by the grams of carbon in the form of plant glucose produced per square metre per year^10, 11^ (Fig. 1), in other words the amount of plant materials in the environment. Low productivity may limit an organism’s ability to withstand physiological stresses. Birds from habitats with fewer plants have lower resistance to oxidative and non-oxidative stress in response to cadmium, hydrogen peroxide, paraquat, tunicamycin, and methane methylsulfonate^12–15^. Additionally, a comparison of 59 temperate bird species from 17 families versus 69 tropical bird species from 29 families shows that species living in lower productivity environments, e.g. temperate birds, have higher metabolic rates than species in higher productivity environments, e.g. tropical birds^12–15^. Higher metabolic rates are linked with increased oxidative stress, immunological stress^16^, and increased cell proliferation, which can lead to cancer^17, 18^ . Still, a recent study suggests that high metabolic rates might be cancer protective^19^. Birds are known to have higher metabolic rates than mammals^20–24^, and lower cancer prevalence than mammals^25–28^.

**Figure 1.**
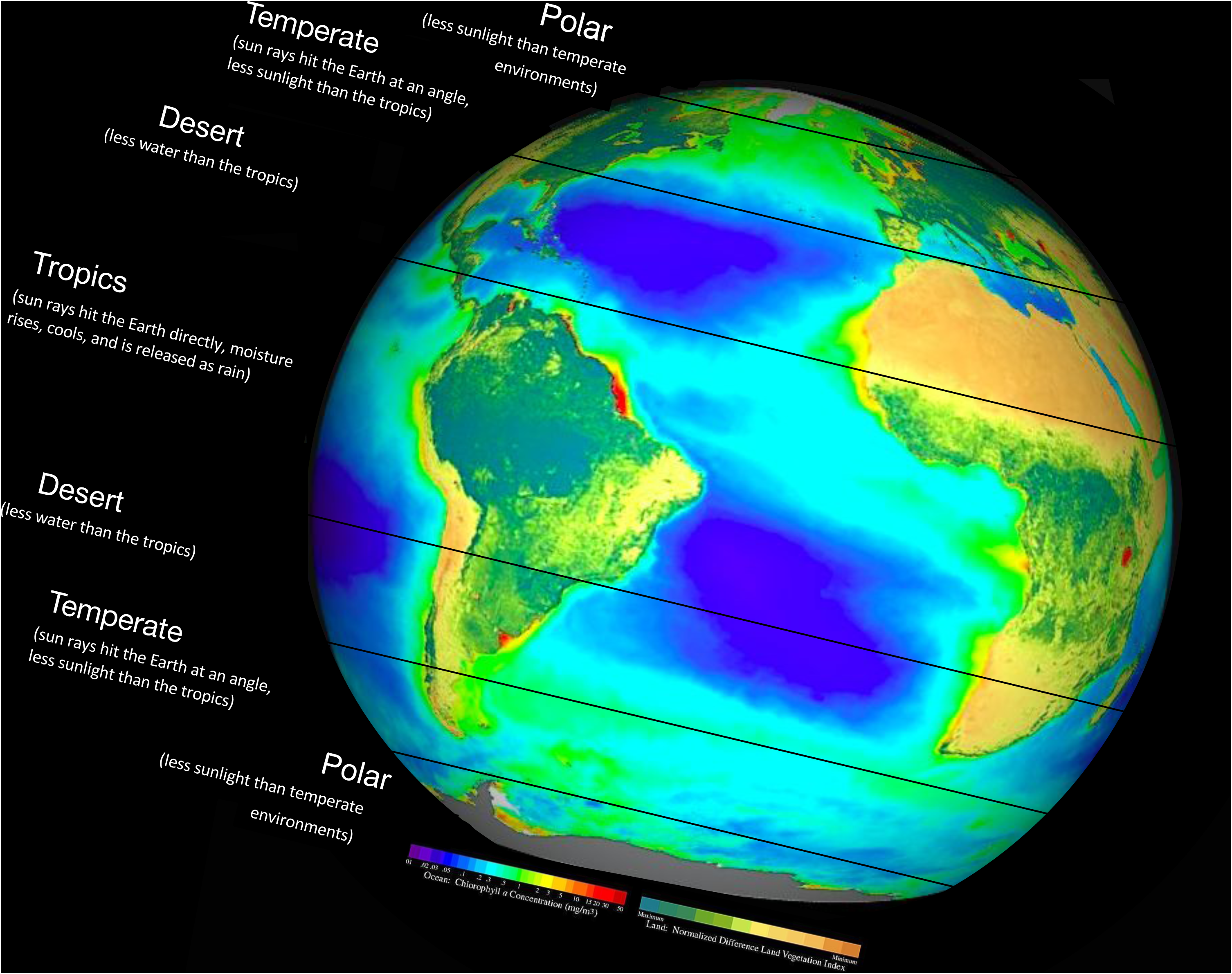
Global distribution of photosynthesis (habitat productivity in grams of carbon produced per square metre per year). The angle of sun rays hitting Earth, the abundance of water, and carbon dioxide, determine the global distribution of photosynthates, oxygen and glucose, through photosynthesis^10, 125, 126^. The green chromophore chlorophyll *a* is involved in this chemical reaction inside the chloroplasts of plants and phytoplankton. We show the global distribution of photosynthesis as concentration gradients of chlorophyll *a*. Dark red and blue-green are areas of high photosynthetic activity in the ocean and on land, respectively. Many grams of carbon in the form of glucose per square metre per year are produced in the tropics [2000 gC/(m² • year)] and freshwaters [2150 gC/(m² • year)], where the photosynthetic reactants sunlight, water, and carbon dioxide, are abundant^127, 128^. Fewer grams of carbon in the form of glucose per square metre per year are produced in deserts [3 gC/(m² • year)], where the photosynthetic reactant water is minimal^10, 125–128^. Reproduced from a satellite image of NASA SeaWiFS Project, Goddard Space Flight Center and ORBIMAGE and https://images.slideplayer.com/25/8083427/slides/slide_27.jpg. Copyright: Wikimedia Commons. We placed the NASA image on a sphere using www.maptoglobe.com to illustrate the part of Earth’s curve closest to the sun, i.e. the tropics.

Lower habitat productivity is also linked with the evolution of meat-eating^29^. Across vertebrates, mammals have the highest cancer prevalence, followed by reptiles, birds, and amphibians^25–28^. Across different orders of mammals, Carnivora have higher neoplasia prevalence than Rodentia, Primates, Artiodactyla, and Diprotodontia^3, 26, 30^, with the latter orders consisting mostly of herbivorous species. Carcinogens are known to concentrate as they move up the food chain, a phenomenon called biomagnification^31–39^ and there is good epidemiological evidence in humans that animal-based diets are more carcinogenic^40, 41^. Higher metabolic rates and larger home ranges (km^2^) have been also observed in species, within the order Carnivora, eating a higher percentage of meat^42^. Therefore, it would be expected that higher trophic levels and species with a larger habitat range would have higher cancer prevalence across vertebrates.

The hypotheses were that: (1) lower habitat productivity; (2) higher trophic levels; (3) larger habitat range; and (4) higher metabolic rates, are associated with higher malignancy and neoplasia prevalence across tissues across species. It was also predicted that the effect of higher trophic levels might be particularly evident in neoplasms of the gastrointestinal tract. These hypotheses were tested on data collected from 244 vertebrate species from zoos, aquariums, veterinary hospitals, and online resources.

## Methods

### Neoplasia Data Collection

We obtained neoplasia prevalence and malignancy prevalence data from 39 institutions on 14,267 individual necropsies across 244 species. Specifically, Mammalia (89 species), Aves (66 species), Reptilia (50 species), Amphibia (24 species), Actinopterygii (13 species), Chondrichthyes (1 species), and Elasmobranchii (1 species) (supplementary data). All animals in the study had been held in human care, including zoos, research facilities, rehabilitation centers or were pets that had been submitted to veterinary clinics. The neoplasia prevalence (including both benign or malignant tumors) or malignancy prevalence was calculated by dividing the number of necropsies that reported neoplasms, or only malignancies respectively, by the total number of necropsies for that species. We calculated tissue-specific neoplasia prevalence or malignancy prevalence by dividing the number of necropsies showing a neoplasm or only malignancy in that tissue by the total number of necropsies for that species. Species were analysed for which there were ≥20 necropsies, as has been done in previous analyses^3^.

### Data Filtering

All infant records were excluded. Infancy was determined if a record’s age was smaller than or equal to that species’ age of infancy (either the weaning age in the life history table, or the average of male and female maturity if there was no weaning age). When there was no recorded age of infancy, the record was considered an infant if it contained any of the following key words: “infant”, “fetus”, “juvenile”, “immature”, “adolescent”, “hatchling”, “subadult”, “neonate”, “placenta”, “newborn”, “offspring”, “fledgling”, “snakelet”, “brood”, “fry”, “fingerling”. Records for which there was no information about the location of the neoplasia and free-ranging animal records were excluded from this study.

For reference, worldwide, adult humans have a cancer rate of approximately 0.18 (https://ourworldindata.org/grapher/share-of-deaths-by-cause), but there is not an estimate of benign neoplasia prevalence in humans, so humans were not included in our database.

### Tissue specific diagnoses

Gastrointestinal malignancy or neoplasia prevalence of each species consists of the malignancy or neoplasia prevalence, respectively, of the oral cavity, esophagus, stomach, gallbladder, bile duct, liver, pancreas, duodenum, small intestine, and colon. A case of a midabdominal mass adhered to a ferret’s stomach and small bowel, and a mass in the pancreatic region of another ferret were included in the gastrointestinal data. The remaining abdominal neoplasms that did not classify as neoplasms of the esophagus, stomach, gallbladder, bile duct, liver, pancreas, duodenum, small intestine, or colon, were included in the non-gastrointestinal neoplasia data.

Specifically, non-gastrointestinal malignancy (or neoplasia) prevalence of each species consists of the malignancy (or neoplasia) prevalence in non-gastrointestinal locations or cell lineage. These locations or cell lineages are the adrenal cortex, adrenal medulla, blood, bone, bone marrow, brain, carotid body, cartilage, dendritic cell, fat, fibrous connective tissue, glandular tissue, glial cell, hair follicle, heart, iris, abdomen, kidney, larynx, lung, lymph nodes, mammary, mast cell, meninges, mesothelium, myxomatous tissue, nerve cell, neuroendocrine tissue, neuroepithelial tissue, nose, notochord, ovary, oviduct, parathyroid gland, peripheral nerve sheath, pigment cell, pituitary gland, neural crest, prostate, pupil, skin, smooth muscle, spinal cord, spleen, striated muscle, synovium, testis, thyroid, trachea, transitional epithelium, uterus, vulva, and when the cancer was considered widespread without an indication of where the primary tumor originated.

### Habitat data collection

To determine the habitat of the 244 species for which there were cancer records, a search was performed predominantly using the Animal Diversity Web (https://animaldiversity.org/) and the International Union for Conservation of Nature (IUCN) red list of threatened species (https://www.iucnredlist.org/). References for the habitat(s) of each species are included in the supplementary data.

Values of habitat productivity for freshwater, tropical, marine, temperate, and desert habitats from previous studies were found^43, 44^ (Table 1A). The productivity of freshwater habitats was classified as the average productivity of swamps, marshes, and river estuaries, whereas the productivity of marine habitats was classified as the average productivity of coral reefs, algal beds, and open oceans (Table 1A). The average habitat productivity of a species was the productivity of its habitat if that species lived in a single habitat, and the average productivity of its habitats if it lived in two or more habitats (Table 1B). Habitats were ordered from lowest to highest productivity (Table 1).

**Table 1A.** Net primary production (in grams of carbon produced per square meter per year) from previous measures of biomass productivity ^43, 44^.

| Producers | Net biomass productivity<br>(gC/(m<sup>2</sup> • year)) = the mass of<br>photosynthate | Natural climate zones of the<br>animals | Average net biomass productivity<br>(gC/(m<sup>2</sup> • year)) in each climate<br>zone |
| --- | --- | --- | --- |
| Swamps and marshes | 2500 | Freshwater (average productivity of<br>swamps and marshes & river<br>estuaries) | 2150 |
| Tropical forests | 2000 | Tropical | 2000 |
| Coral reefs | 2000 | Marine (average productivity of coral<br>reefs, algal beds, and open oceans) | 1375 |
| Algal beds | 2000 |  |  |
| River estuaries | 1800 |  |  |
| Temperate forests | 1250 | Temperate | 1250 |
| Tundra | 140 |  |  |
| Open ocean | 125 |  |  |
| Deserts | 3 | Desert | 3 |

**Table 1B.**
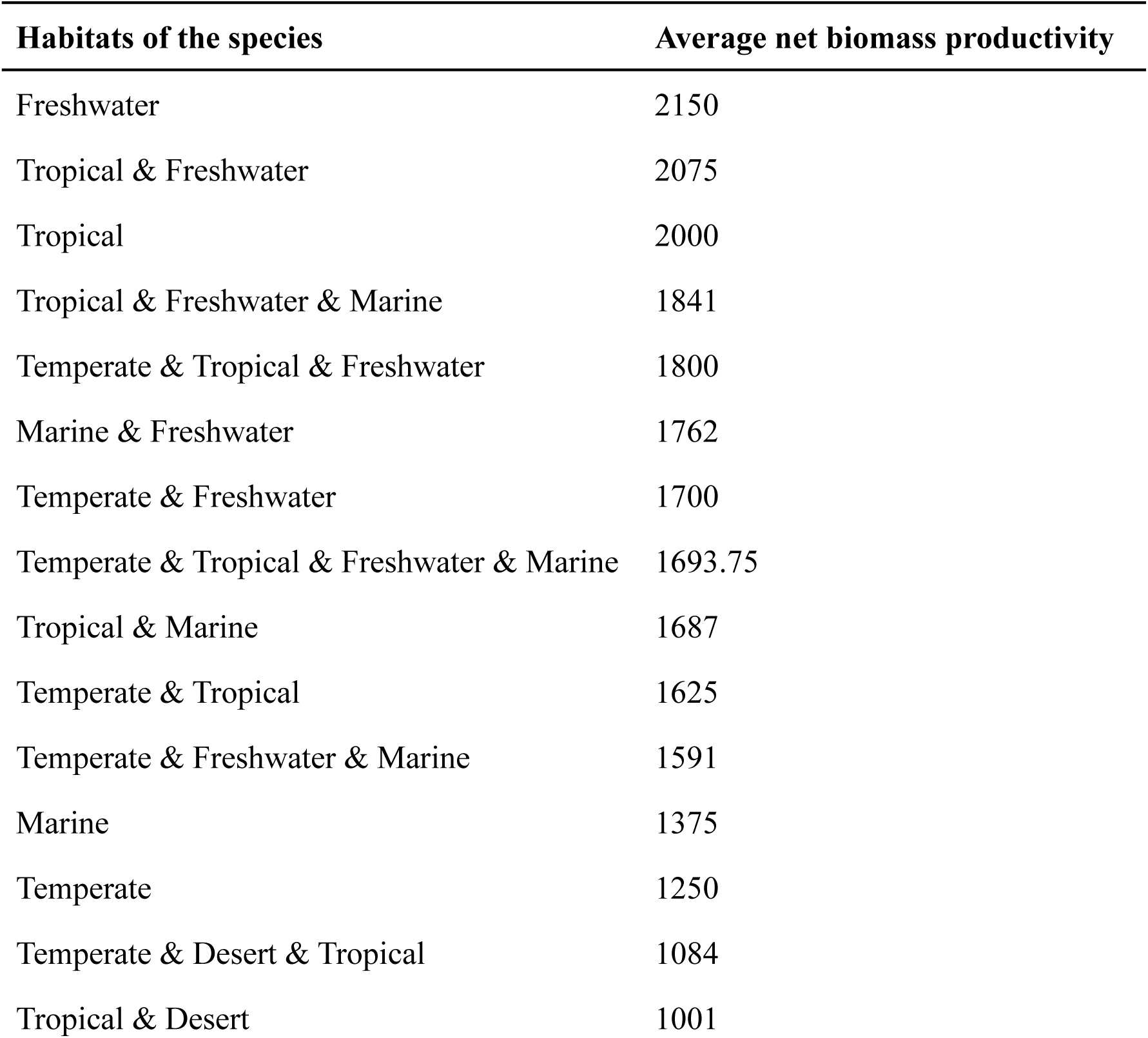
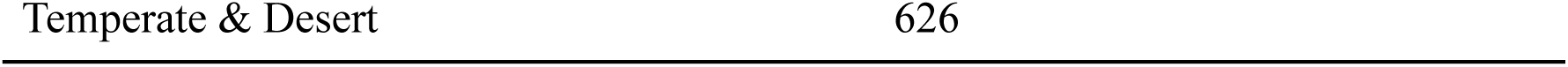
Average net biomass productivity, measured in grams of carbon produced per square meter per year, of the habitats in our database. Rows with multiple habitats are averaged values of these habitats from Table 1A.

To find how many habitats each species lives in, we used the habitat information (supplementary data; Table 1), and categorised each species by their number of habitats (*one*, *two* or *three or more* habitats). Few species lived in four or five different habitats, so these these species were classified as living in three or more habitats.

### Diet data collection

When considering the effect of diet on neoplasia prevalence, there are potentially two different effects: the diet that the species has evolved to eat and the diet those animals are actually fed under human care as pets or in zoological institutions. The dietary data for the animals in human care are not available for this study.

To find the natural diet of each of the 244 species in our database in the wild, a search was performed in the Animal Diversity Web (https://animaldiversity.org/), the IUCN red list of threatened species (https://www.iucnredlist.org/), and other resources^45–50^. Each ecosystem can have many trophic levels. For example, marine ecosystems usually have more trophic levels than terrestrial ecosystems^51–53^. Each organism may eat many types of food creating complex interactions that appear as a food web^54^. In order to test hypotheses in all ecosystems, based on the trophic pyramid^55^, and the food pyramid of low to high cancer incidence in humans^40, 41^, each species was classified according to their primary diet into four trophic levels: herbivores, invertivores, primary carnivores, and secondary carnivores. Decomposers, such as certain bacteria and fungi, were not included in these analyses, because there are not malignancy or neoplasia prevalence data for these species. In these classifications, herbivores include species eating primarily plants (fruit, seeds, bark, leaves, flowers, roots) and/or fungi. Invertivores include species eating primarily invertebrate animals such as arthropods, cnidarians, echinoderms, comb jellies, annelids, and/or molluscs. Eating primarily vertebrates automatically assigned a species as a primary carnivore. To verify this animals were evaluated if their prey were primary carnivores. If the prey were also primary carnivores, these species (i.e. the predators of primary carnivores) were classified as secondary carnivores (Supplementary Data). The diet and habitat of a species may change over time. In this paper, species were analysed according to their primary diet in the wild as reported in recent years. If data on the diet of juveniles in the wild were available, we classified these species according to the diet of adults in those species.

### Inferred metabolic rates

To find the metabolic rate of each species we used the following formula M ^−1/4^ • e^−E/kT^. This formula is part of a more extended formula for metabolic rates “B = bo • M ^−1/4^ • e^−E/kT^” (where B = mass-specific metabolic rate; bo = a coefficient independent of body size and temperature; M = adult mass in grams, E = 0.65 eV, k = 8.62 • 10^−5^ eVK^−1^, T = body temperature in Kelvin)^56–59^. Since there is not a known value for bo in most species, we measured M ^−1/4^ • e ^−E/kT^ for every species. Adult mass data was obtained for the species where their weight was known^60, 61^ (AnAge Database https://genomics.senescence.info/species/index.html). The average body temperatures were found for 81 out of the 244 species in our study. Most of these average body temperatures were from Moreira et al.’s recent study^62^ on 1,721 species of land vertebrates, including amphibians, mammals, reptiles, and birds collected from the lab, field, and an ‘unclear’ location. Body temperature for the 15 fish species was unknown and not included in any analyses. Average body temperature of marine mammals was generated from van Wijngaarden et al, Mense et al., and McCafferty et al.^63–65^. The mass-specific metabolic rate was estimated as the unweighted average of body temperature. In order to use these body temperature values in “M ^−1/4^ • e ^−E/kT^”, the body temperatures was converted from Celsius to Kelvin.

### Statistical Analyses

All were performed analyses in R version 4.0.5^66^ using the R packages CAPER^67^, phytools^68^, geiger^69^, tidyverse^70^, and powerAnalysis (https://github.com/cran/powerAnalysis), and performed Phylogenetic Generalised Least Squares (PGLS) regressions to take into account the phylogenetic non-independence among species. To perform a PGLS regression, a phylogenetic tree (phyl file) was made using the NCBI Tree creator (https://www.ncbi.nlm.nih.gov/Taxonomy/CommonTree/wwwcmt.cgi). PGLS analyses have also been previously used to compare neoplasia prevalence and life history variables across mammals^1^. In the analyses where the dependent variable was malignancy or neoplasia prevalence, we weighted the analyses by 1/(square root of the number of necropsies per species) (from Revell ^68^). A Bonferroni corrections was performed to adjust for multiple testing.

The trophic level, average habitat, minimum habitat productivity, and number of habitats were used as categorical variables. Whereas the malignancy prevalence, neoplasia prevalence, average body temperature, “M ^−1/4^ • e ^−E/kT^”, and average habitat productivity (Table 1), were used as numerical variables. The average body temperatures were transformed to the power of 2, and the value of “M ^−1/4^ • e ^−E/kT^” to the power of 0.1 to normalise the distribution of data.

## Results

This study found that species from lower productivity habitats have more malignancies, more tumors, more gastrointestinal cancer and gastrointestinal neoplasms in general (Table 2). These associations were examined in two ways. First, using the minimum productivity habitat occupied by a species and controlling for the species position in the trophic pyramid (Fig. 2). Second, using the average productivity of the habitat(s) a species occupies and controlling for the higher levels of neoplasms in mammals^25^ (Fig. 3). These correlations, however, between habitat productivity and malignancy or neoplasia prevalence were not significant after applying Bonferroni corrections for multiple testing (Table 2).

**Figure 2.**
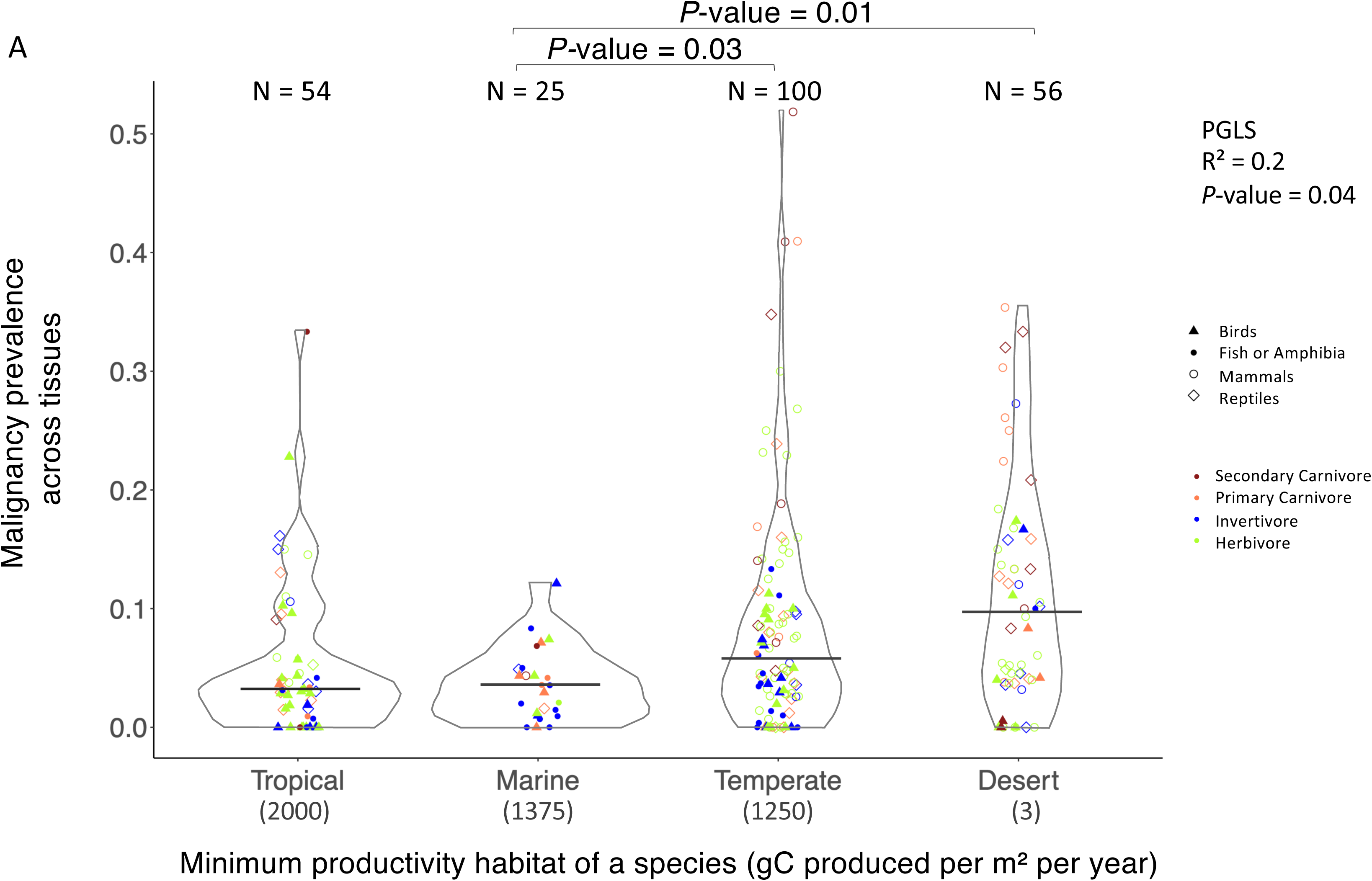
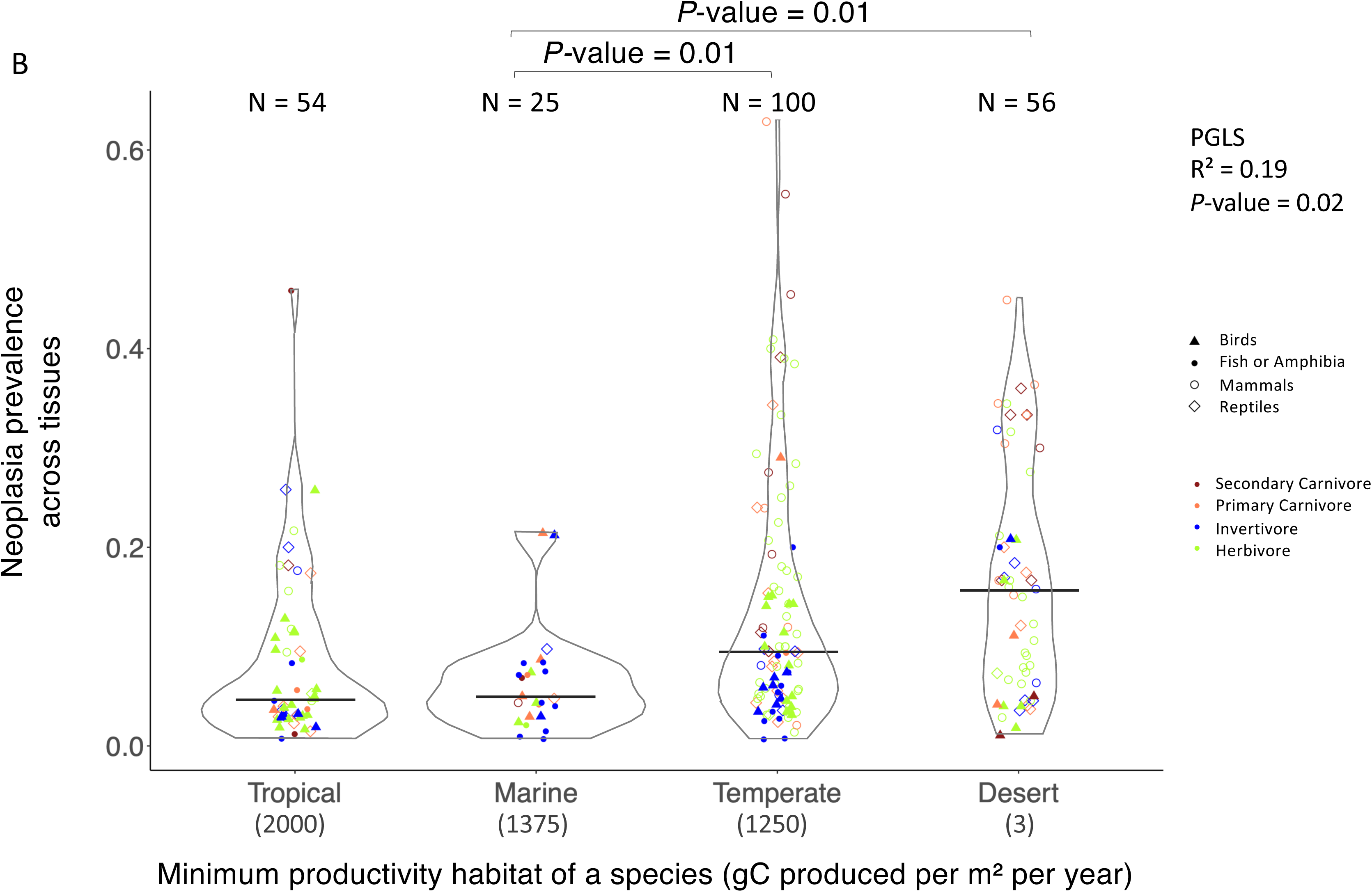
Malignancy and neoplasia prevalence across habitats. Malignancies (A) and neoplasias (B) are more prevalent in species that live in deserts. Productivity is measured in grams of carbon produced per square meter per year. Each species is classified according to its minimum productivity habitat among the habitats in its range. Habitats are ordered from left to right according to their productivity, from high to low productivity. Each dot is a species, and each category on the x axis consists of ≥10 species. Colours show the trophic levels of each species, and the shapes indicate the major clades. The number of species (N) in a category is noted above each category. The horizontal black line in each category shows the median malignancy or neoplasia prevalence in that category. *P-*values between categories that are significantly different are shown. All *P-*values are provided in the supplementary data and Table 2.

**Figure 3.**
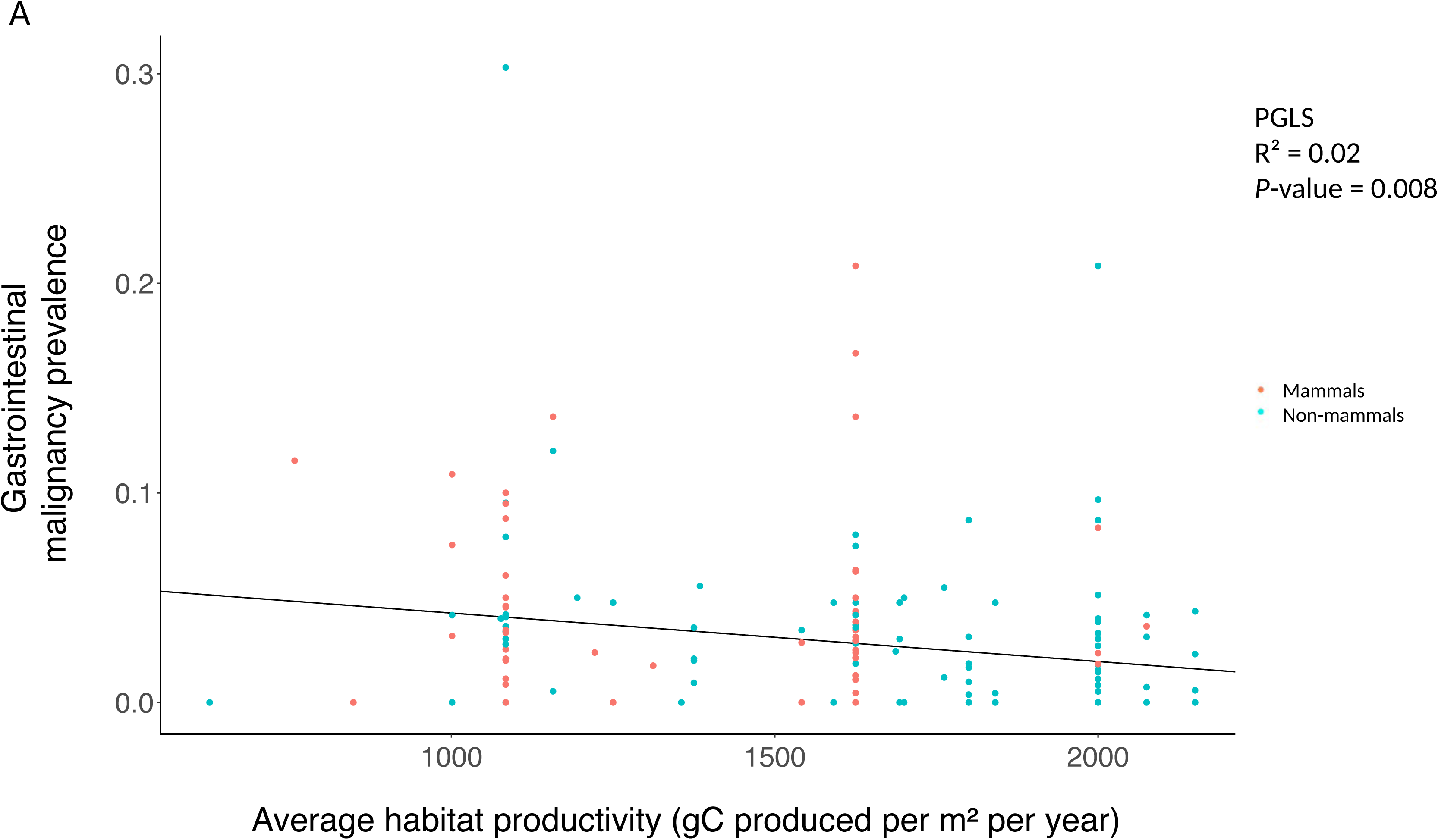
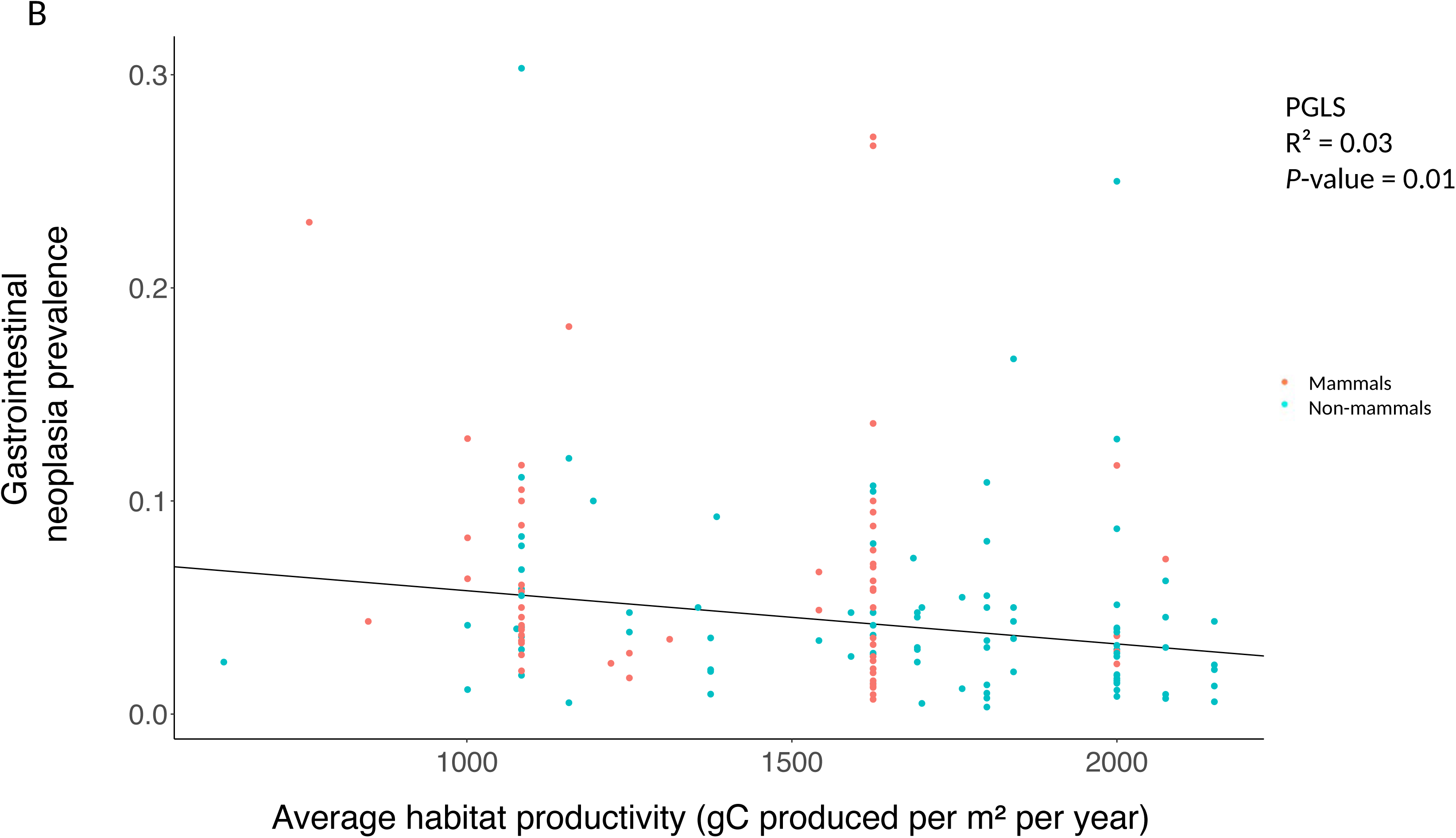
Gastrointestinal malignancy (A) and neoplasia prevalence (B) are higher in species living in habitats which on average have low productivity. Habitat productivity is measured in grams of carbon produced per square metre per year. Each dot is a species. See Table 2 for statistical analyses.

**Table 2.**
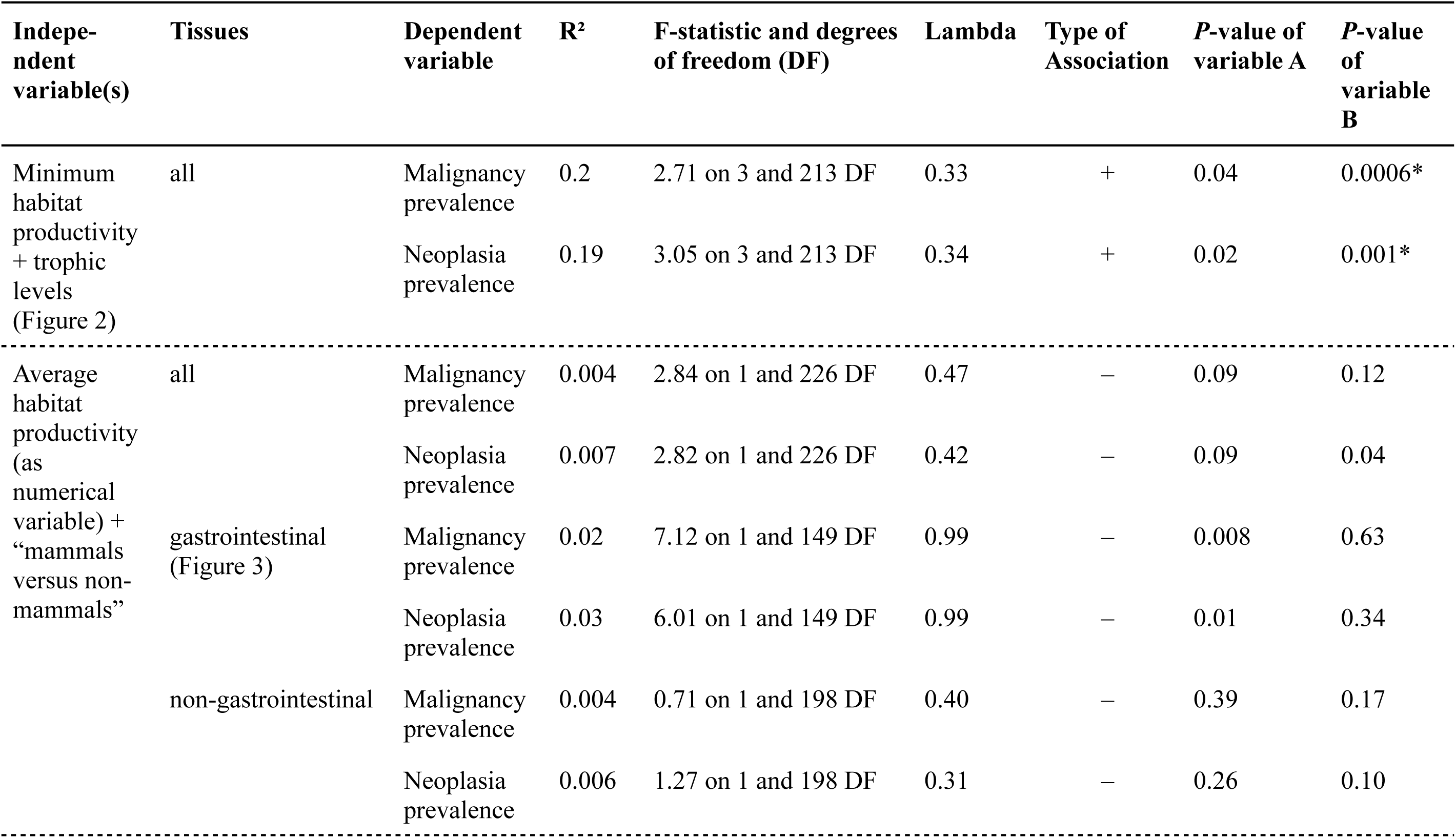
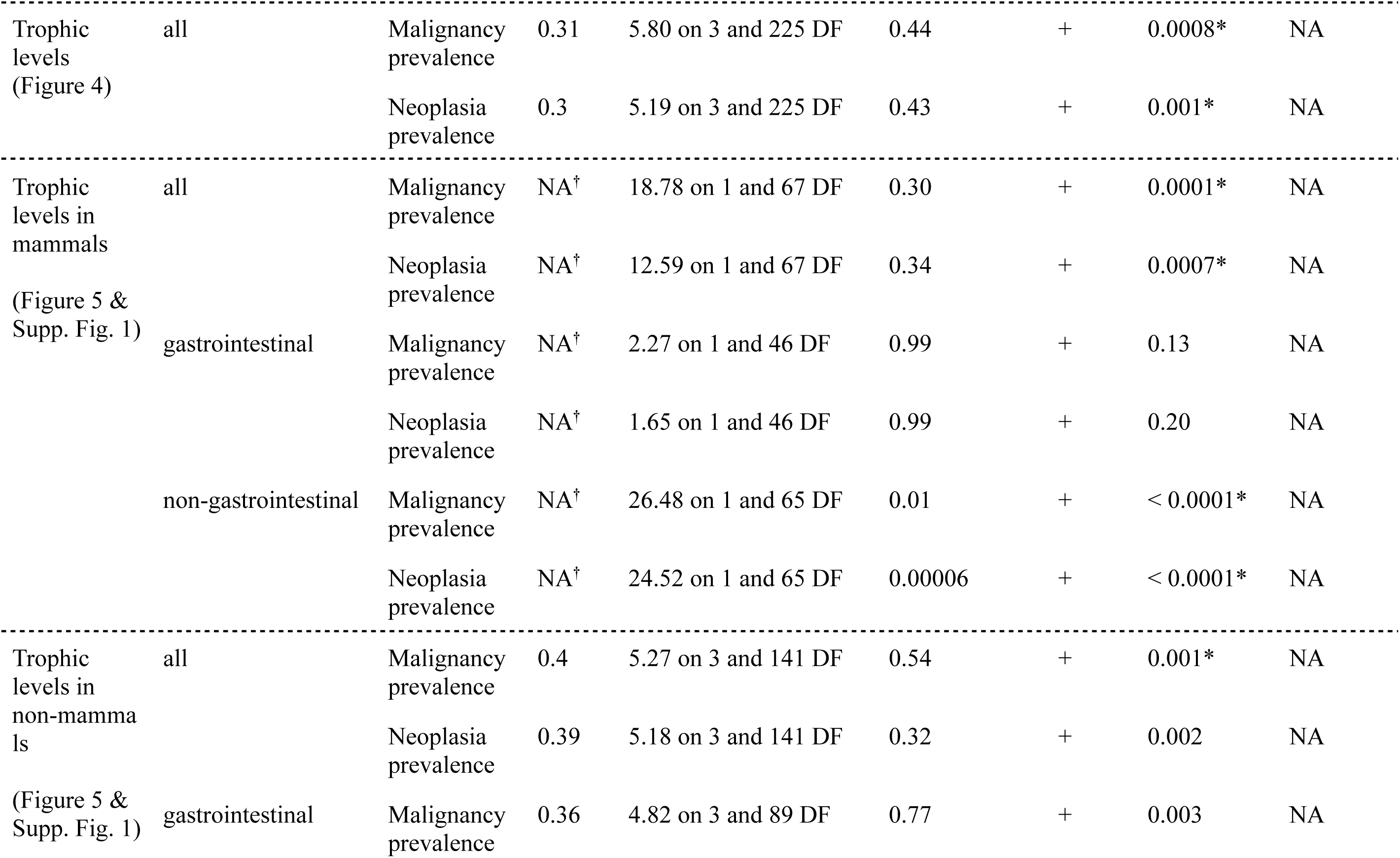
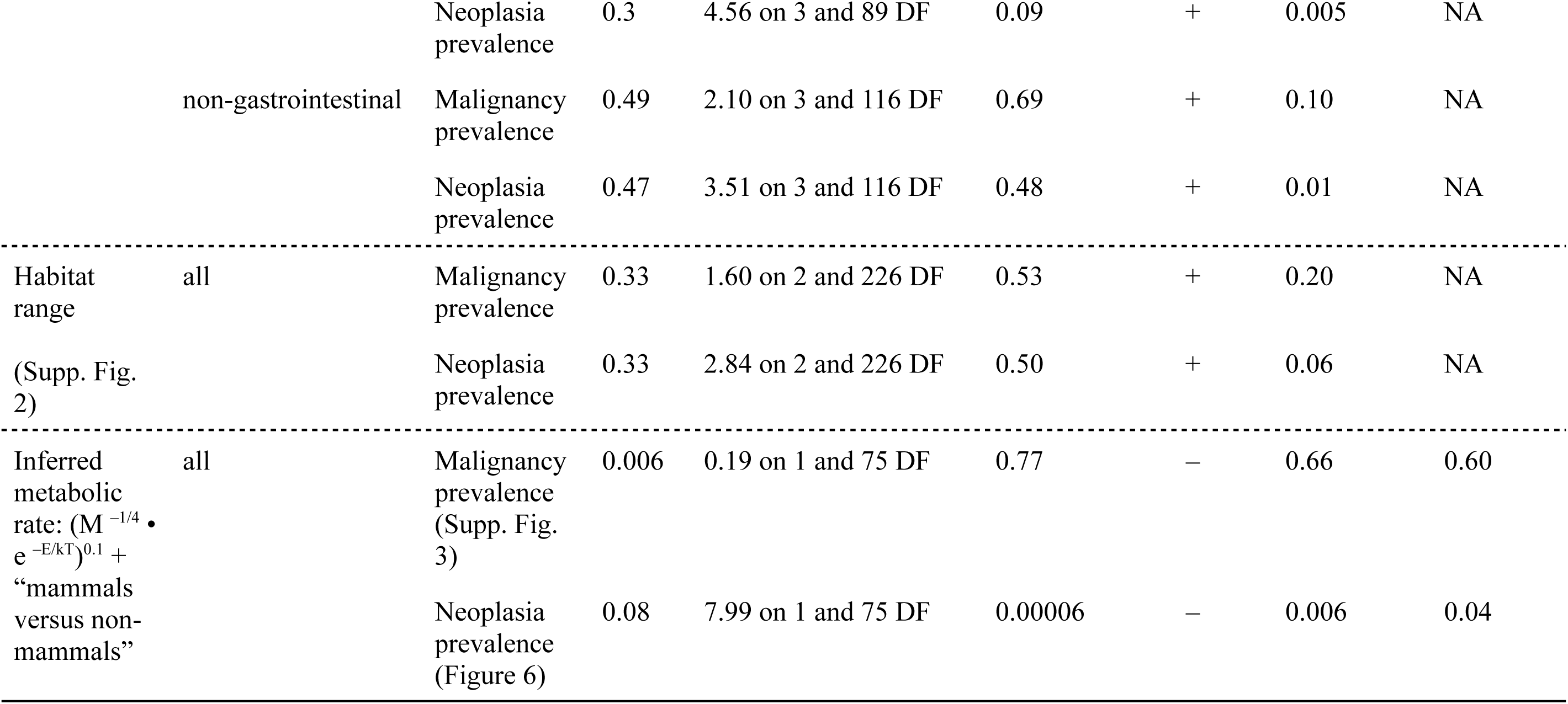
Phylogenetic generalised least squares (PGLS) regression summary results of the figures in the main text and in the supplementary materials. High values of lambda indicate that the signals can be mainly explained by common ancestry among species. With 26 tests, the most conservative Bonferroni correction requires a *P*-value < 0.0019 in order to be considered statistically significant, which are highlighted with a * next to the *P*-value. In the 1st P-value column the *P*-value are reported for the first variable (which we call variable A in the multivariate analysis), in the 2nd *P*-value column the *P*-value is reported for variable B, in the F-statistics column we report the F-statistics of variable A, and in the “Type of Association” column we report whether there is a positive (+) or negative (-) association between the independent variable A and neoplasm or malignancy prevalence. If the independent variable is categorical, the sign (+ or –) of the majority of between-group comparisons is reported. ^†^ indicates that the R² value was not available given that the categorical independent variable only included two groups (herbivores and primary carnivores).

There were no significant associations overall between habitat range and malignancy prevalence across tissues or neoplasia prevalence across tissues without controlling for increased cancer in mammals, and after applying a Bonferroni correction (Supp. Fig. 2). There is evidence of a trend (*P*-value = 0.01), however, in species living in three or more habitats having higher neoplasia prevalence than species living in one habitat (Supp. Fig. 2B).

Higher trophic levels tend to have higher malignancy and neoplasia prevalence across tissues (Fig. 4), both within mammals and within non-mammals (Fig. 5 & Supp. Fig. 1; Table 2). Also, in mammals, higher trophic levels have higher non-gastrointestinal malignancy and neoplasia prevalence. Gastrointestinal neoplasia prevalence in mammals seems to largely be determined by phylogenetic closeness (Lambda = 0.99), potentially between the species in the order Carnivora relative to the other mammalian orders, and there was little evidence that gastrointestinal neoplasia prevalence was influenced by trophic level (Fig. 5 & Supp. Fig. 1; Table 2). In contrast, in non-mammals, higher trophic levels had higher non-gastrointestinal neoplasia prevalence as well as higher gastrointestinal malignancy and neoplasia prevalence (Fig. 5 & Supp. Fig. 1; Table 2). Most of these relationships remained statistically significant after a Bonferonni multiple testing correction (Table 2).

**Figure 4.**
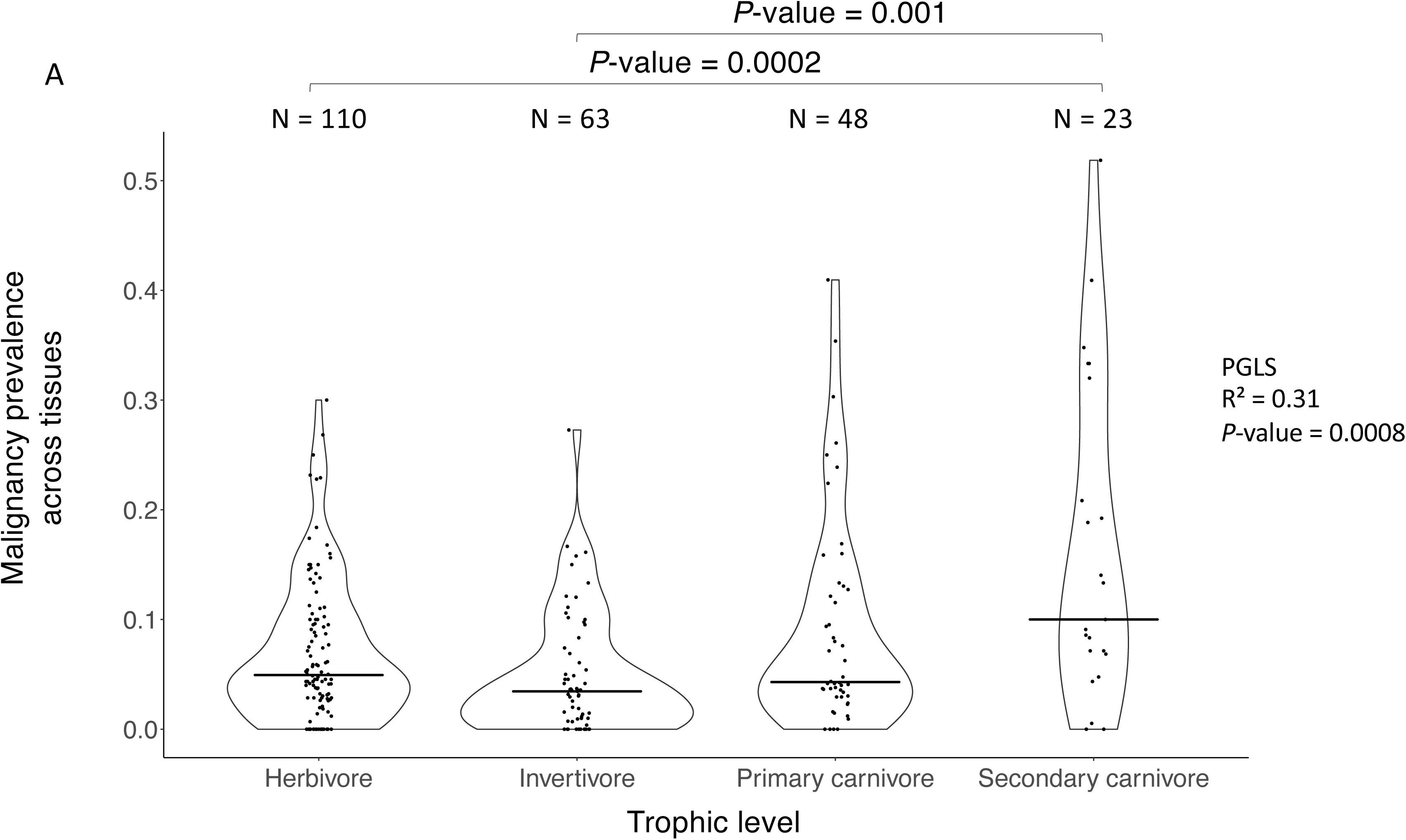
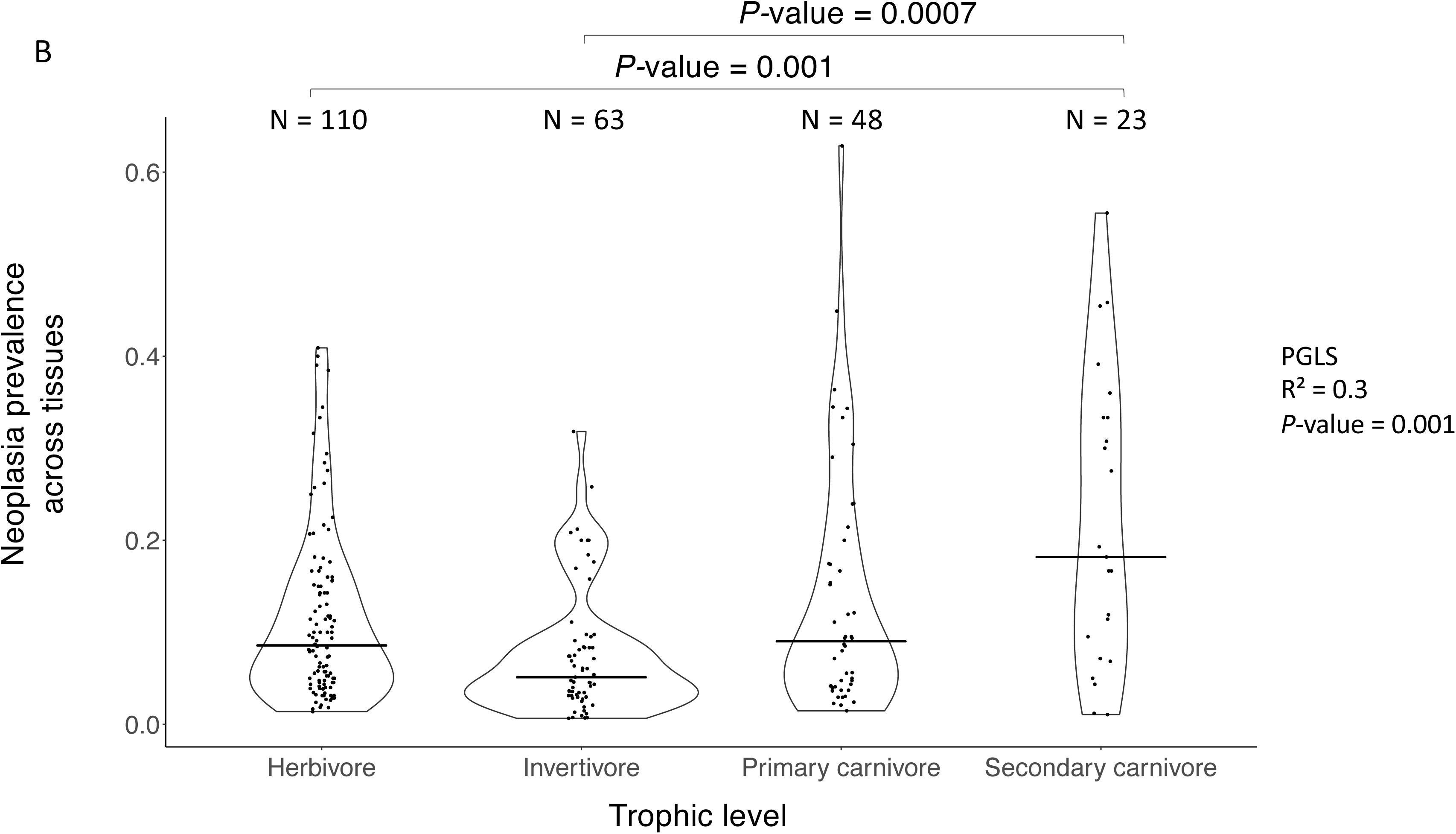
Malignancy (A) and neoplasia (B) prevalences are significantly higher in higher trophic levels. Each dot is a species, and each trophic level consists of ≥10 species. The number (N) of species in each trophic level is indicated above each trophic level. The horizontal black line in each category shows the median malignancy or neoplasia prevalence in that category. *P*-values are provided between categories that are significantly different, and for all comparisons in the supplementary data and Table 2.

**Figure 5.**
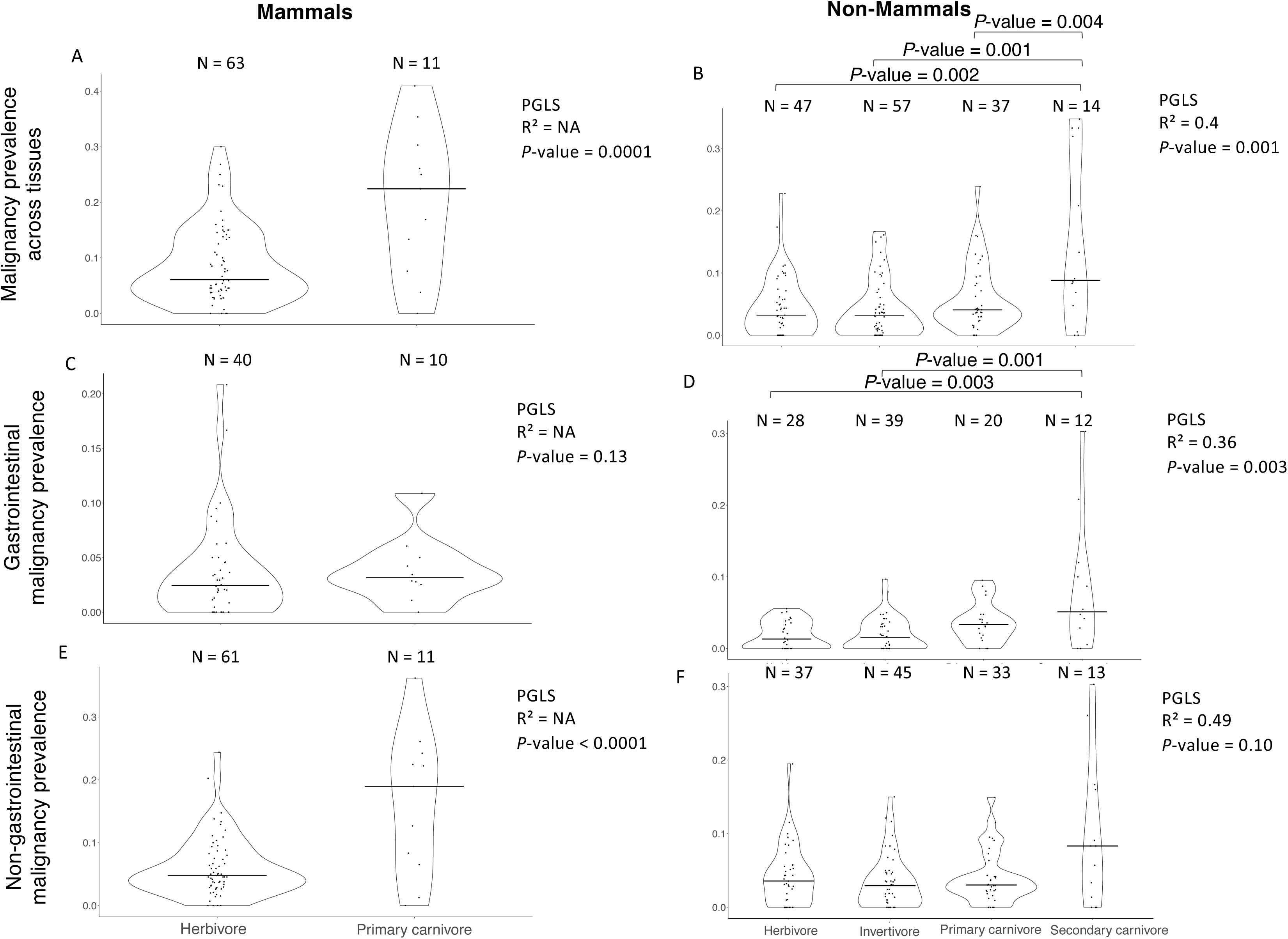
Malignancy and neoplasia prevalence in relation to trophic levels in mammals versus non-mammals. Panels A-F show malignancy prevalence as a function of trophic levels in mammals versus non-mammals, for all tissues (Panels A & B), gastrointestinal malignancies (Panels C & D), and non-gastrointestinal malignancies (Panels E & F). Malignancy prevalence is always higher in higher trophic levels except for gastrointestinal malignancy prevalence in mammals (Panel C) and non-gastrointestinal malignancy prevalence in non-mammals (Panel F). Each dot is a species, and each trophic level consists of ≥10 species. The number (N) of species in each trophic level is indicated above each trophic level. The horizontal black line in each category shows the median malignancy prevalence in that category. *P*-values are provided between categories that are significantly different, and for all comparisons in the supplementary data and Table 2.

Species with lower inferred metabolic rates [measured as (M ^−1/4^ • e ^−E/kT^)^0.1^, where M = adult weight, and T = average body temperature] had higher neoplasia prevalence (but not higher malignancy prevalence) across tissues when controlling for increased cancer in mammals (Fig. 6; Table 2, and Supp. Fig. 3), however, this association was not statistically significant after applying Bonferroni corrections for multiple testing (Table 2).

**Figure 6.**
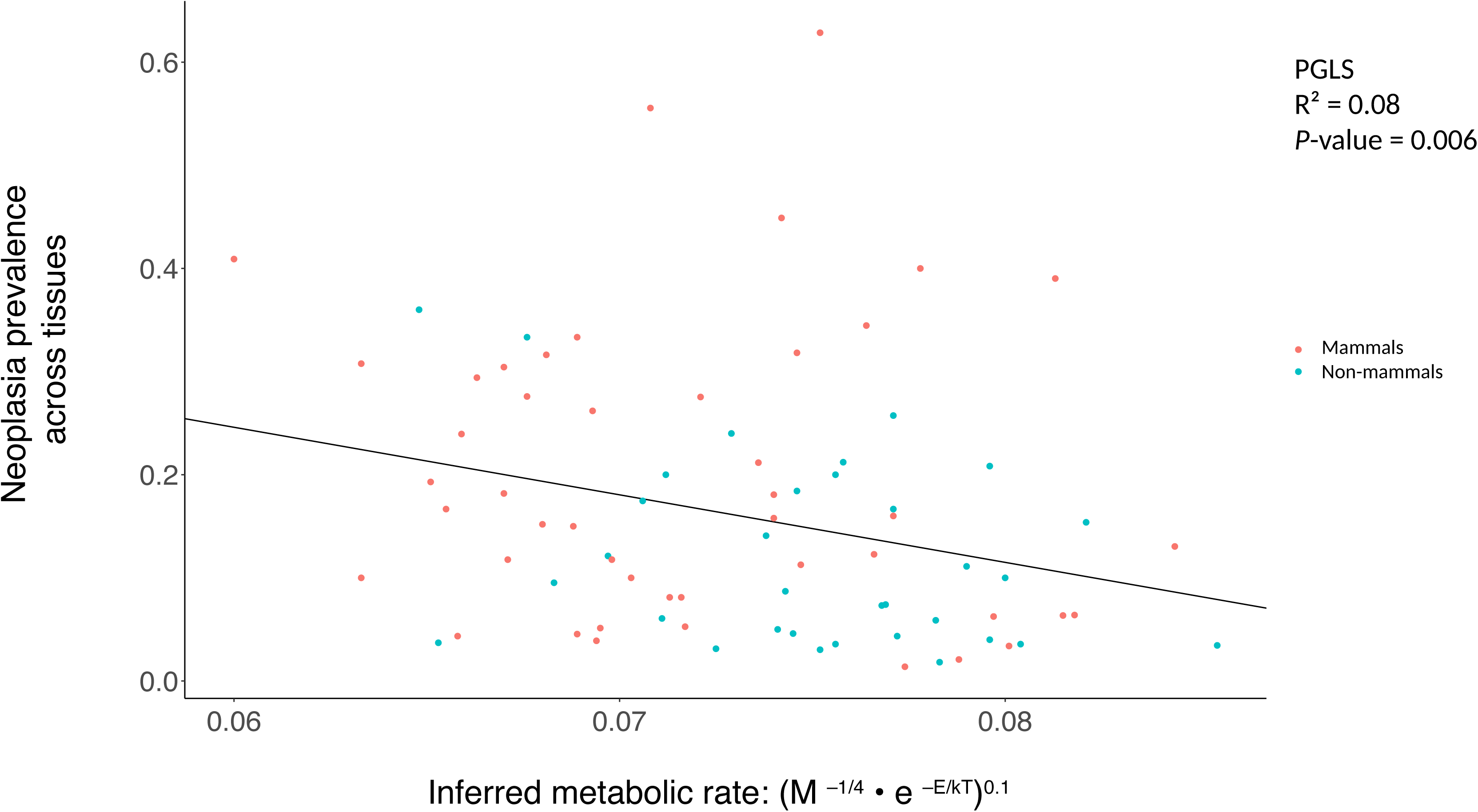
Neoplasia prevalence across tissues is higher in species with lower inferred metabolic rates. The formula M^−1/4^ • e ^−E/kT^ multiplied by *bo* is a measure of metabolic rate for each species. We do not know the *bo* value [*bo* = a coefficient independent of body size and temperature from ^58^] for each species, so we measured the M^−1/4^ • e ^−E/kT^ values for each species (where M = mass, E = 0.65 eV, k = 8.62 • 10^−5^ eVK^−1^, and T = temperature in Kelvin)^56^. Each dot is a species.

Average body temperature was not significantly correlated with malignancy or neoplasia prevalence across tissues without controlling for increased cancer in mammals (Supp. Fig. 4). Also, inferred metabolic rates (M ^−1/4^ • e ^−E/kT^)^0.1^ were lower in primary carnivores than invertivores (Supp. Fig. 5).

The majority of species in lower trophic levels in our database lived in higher productivity habitats, whereas the majority of species in higher trophic levels lived in lower productivity habitats (Supp. Fig. 7). When controlled for trophic level, there was still evidence that low productivity habitats are still associated with more neoplasia (P=0.02) and malignancy (P=0.04), though neither was significant after Bonferroni correction for multiple testing.

## Discussion

Cancer prevalence varies across species, for mostly unknown reasons^3, 25, 26, 30^. This study hypothesised that ecological variables would explain some of this variance in cancer prevalence. Evidence was found that malignancy and neoplasia prevalence are higher in species from lower productivity habitats (Table 2). There was also evidence that neoplasia prevalence is higher in species with lower inferred metabolic rates (Table 2).

The strongest effect found was that malignancy and neoplasia prevalence are higher in higher trophic levels, particularly in secondary carnivores (Figs. 4 & 5, Table 2). A recent study in mammals on a different dataset showed an association between cancer mortality risk and animal content in a species’ diet^3^. Here this association was expanded across vertebrates, and discovered that this was partly driven by higher levels of tumors, both benign and malignant, in vertebrate species that are secondary carnivores. It is possible that some of the association between trophic level and cancer prevalence was due to biomagnification of non-degradable toxic chemicals, such as heavy metals, pollutants, microplastics, and pesticides^71^, causing detrimental health effects as their concentration increases in animal tissue at every step higher in the trophic pyramid^31–37, 39^. This study also discovered strikingly low rates of neoplasia prevalence in species that mainly feed on insects or other invertebrates (Figs. 4 & 5, Supp. Fig. 1). Trophic levels explain much of the variation in malignancy and neoplasia prevalence across tissues across species in our database (R² as high as 0.31; Table 2).

Caution should be used when applying observations across species to within species; however, these observations of increased cancer prevalence at higher trophic levels were consistent with many studies in humans showing an association between cancer and consumption of animal products^72–77^. In humans, consuming products low in photosynthates, such as red meat, animal fat, and/or processed food, leads to the production of N-nitroso compounds, secondary bile acids, hydrogen sulfide, and reactive oxygen species^41, 78^. These chemicals increase DNA damage and substitution rates, reduce mucus production, degrade the extracellular matrix, cause inflammation and tumor proliferation^40, 41, 78–82^. Products abundant in photosynthates, such as fruit and vegetables^83–90^, produce more vitamins^41^, antioxidants^41^, flavonoids^41, 91^, and glucosinolates^41^, which reduce DNA damage and inflammation^41^, degrade toxins, limit pathogen growth^41^, tighten cellular junctions^41^, maintain the barrier between intestinal cells^41^, and inhibit tumor growth^40, 41, 79, 92^. Perhaps it is no coincidence that the majority of already known cancer-resistant mammals, such as naked-mole rats, blind mole rats, and elephants, are herbivores^93–95^. Despite these observations, the story is likely to be more complex. It is likely that species with meat-based diets have evolved specialised adaptations to consuming meat that may mitigate the negative effects of carnivory.

Invertivores in most of these analyses had the lowest median cancer prevalence and neoplasia prevalence of all trophic levels (Fig. 4 & 5; Supp. Fig. 1). Previous studies have shown that insectivorous fish have higher mean concentrations of total mercury in their tissues than herbivorous fish^96^. The majority of herbivores in our database, however, were endotherms, whereas the majority of invertivores in our database were ectotherms. Ectotherms are known to have lower concentrations of persistent organic pollutants in their body than endotherms, because endotherms of the same size consume relatively more food as a means of maintaining body temperature. Therefore, this lower concentration of persistent organic pollutants in the body of ectotherms^97, 98^, in combination with the fact that there are more ectotherms than endotherms in our invertivore trophic level, may explain the lower cancer prevalence in invertivores than herbivores (Fig. 4 & 5; Supp. Fig. 1). This study found, however, that cancer prevalence was not significantly different between endotherms and ectotherms when controlling for trophic levels (multivariate PGLS, *P*-value = 0.46; Lambda: 0.42; R²: 0.05). This lower cancer prevalence in invertivores may also be due to the presence of insect chemicals, such as chitin, in their diet. Chitin is a polymer of *N-*acetylglucosamine, a derivative of glucose, and is the primary substance of the exoskeleton of almost all invertebrates and most fungi. There is evidence that chitin and many other insect-derived compounds have antioxidant, anti-inflammatory, and anti-cancerous properties^99–101^.

Species at high trophic levels tend to live in many habitats^42, 102, 103^ (Supp. Fig. 6) and in low productivity habitats in particular (Supp. Fig. 7). This larger habitat range of carnivores versus herbivores, may be explained by unpredictability in the abundance and wide dispersal of their prey. Carnivores have relatively more mobile prey than herbivores, and this possibly led to adaptations related to movement in a broader habitat range in carnivores than herbivores ^104^. Species living in larger habitat ranges, however, do not have higher malignancy or neoplasia prevalence across tissues than species living in one habitat (Table 2).

DNA base pair substitution (mutation where a pair of bases changes in the DNA) rate is sometimes modelled as a function of the inferred metabolic rate^56–59, 105^, and so metabolic rate may be associated with cancer due to its relationship to mutation rate. Additionally, higher metabolic rates are characteristic of higher trophic levels^42, 106, 107^. Still, birds have higher metabolic rates than mammals^20–24^, and lower cancer prevalence than mammals^25–28^. This study found that inferred metabolic rates did not differ significantly between trophic levels (Supp. Fig. 5), and inferred metabolic rates showed a negative correlation with neoplasia prevalence across tissues (Figure 6; *P*-value = 0.006) which was not significant after applying Bonferroni corrections (Table 2). This discrepancy with Muñoz-Garcia et al.’s findings may be due to the fact that this study measured inferred metabolic rates (M^−1/4^ • e^−E/kT^ from the metabolic theory of ecology, where M is adult weight and T is their body temperature)^56, 57^ across the tree of life, whereas Muñoz-Garcia et al.^42, 106, 107^ measured the basal metabolic rate or rate of oxygen consumption per species only within the order Carnivora.

Why haven’t species at lower productivity habitats, higher trophic levels, or with lower metabolic rates evolved better cancer defences than species in higher productivity habitats, lower trophic levels, or higher metabolic rates? Resource scarcity in lower productivity habitats and bioaccumulated toxins in higher trophic levels may limit adaptive responses to cancer defences. Imagine a damaged piece of DNA in two different environments: tropics and deserts. In the tropics, the resources for fixing/maintaining^108, 109^ the broken DNA are more likely to be available than in the deserts. Some tropical birds with slower life histories are more resistant to oxidative stress than temperate birds with faster life histories^12–14, 110^. Tropical predators may also have more resources for detoxifying the bioaccumulated toxins. In other words, tropical birds may invest more resources in cellular maintenance, such as cancer protection, than temperate species^12, 111^. Bioaccumulated natural (and artificial) toxins in high trophic levels, and loss of heat, waste, and dead matter in every trophic level, can reduce population size in high trophic levels and thus limit adaptive responses to future environmental stressors^112^. Therefore, even though damages in the DNA can potentially happen in many different environments and trophic levels, their chance of repair may be higher in species of higher productivity environments and lower trophic levels.

### Limitations & future directions

There were 244 species in this database, however, some data were missing or were not precise for many of those species. For example, the adult weight and body temperatures of 188 and 81 species, respectively, were available. This limits statistical power for identifying relationships between variables and cancer prevalence. Some taxa, such as birds, have too few species per habitat or trophic level (e.g. only 8 primary carnivores and only 2 secondary carnivores within birds) to identify associations between habitat or trophic level and cancer prevalence just within that taxon. Another caveat in our study is that precise data on the productivity of the specific habitats inhabited by each species does not exist, nor are there data on the amount of a species range that is covered by each habitat. The number of grams of carbon produced per square metre per year is an average over the whole year based on the habitat of each species^43, 44^ (Table 1A & 1B; supplementary data). Furthermore, although the outcomes of malignancy and neoplasia prevalence in this study are based on a minimum of 20 necropsies per species, future studies with additional individuals may find other relationships than are currently available in this dataset.

The estimates of metabolism are inferred rather than directly measured (Methods). The *bo* value, a coefficient independent of body size and temperature in the metabolic rate equation, is unknown for most of these species (Supplementary Data). Ideally, future studies will provide more accurate measurements of each species’ average metabolic rate and the geographical coordinates of wild animals.

This database is composed of animal populations housed under human care in zoos, aquariums, and as pets submitted to veterinary practices (Supplementary Data). In a study across over 50 mammalian species, all Carnivora lived longer in managed populations than in free-ranging wildlife populations, whereas not all species from mostly herbivorous orders had an extension of lifespan in human managed populations^113^. Cancer is a disease of aging, and this enhanced longevity in species at higher trophic levels may expose them to developing more cancer. So the fact that high trophic levels have higher cancer prevalence than low trophic levels in these data suggests that this may not be as strong an effect in wild populations. There may be mismatches in the food given to each animal in different zoos due to the different location and availability of food in each zoo; however, some of the institutions in this study that belong to the Association of Zoos and Aquariums (AZA) accredited institutions follow similar feeding guidelines. There may also be mismatches in the habitat and diet of animals from various institutions in our database, compared to animals living in the wild, that may affect their susceptibility to cancer. For example, in AZA accredited institutions (feeding guidelines: https://www.aza.org/animal-care-manuals and https://nagonline.net/tag/animal-care-manuals/), lions eat predominantly raw meat or cat food; European polecats eat primarily cat food supplemented with fruit and vegetables or

Mazuri® Ferret Diet; meerkats eat primarily cat food, Royal Canine® Vet Diet, Natural Balance® Carnivore 10% food, or fruit and vegetables; and North American river otters eat primarily cat food, horsemeat, or beef-based diets supplemented with vegetables, minerals, vitamins, and raw fish. Whereas in the wild, meerkats are invertivores and North American river otters are secondary carnivores (Supplementary Data). The diets that were fed to the specific individuals in our database are not available. Also, species in human care may experience different climate conditions than their environment in the wild. For example, species may live in a desert-like environment under human care, but in a temperate environment in the wild. Future studies that track the diet of individuals worldwide in zoos, aquariums and veterinary hospitals, may determine if diets and habitats are better or worse predictors of cancer prevalence compared to the experienced habitats and diets of the individual animals being studied. This study was based on the primary diet and habitat the species occupied in the wild which may have shaped the evolution of each species, its cancer prevalence and cancer defences over thousands of years.

We tested the effect of environmental factors such as habitat productivity, habitat range, and diet, on malignancy and neoplasia prevalence across the tree of life. There may be additional factors, however, within these or neighboring habitats that affect malignancy and neoplasia prevalence across species, that were not tested. The use of hormonal contraceptives in human managed populations may contribute to higher cancer prevalence in some of our species, particularly in females. When testing for a sex bias in cancer risk in the order Carnivora, however, Vincze et al.^3^ did not find any significant sex bias in cancer risk, indicating that hormonal contraceptives do not explain the increased cancer prevalence in the populations of Carnivora under human care that were examined in that study. A study from 1977 found that the majority of neoplasms in mammals were found in the lung^25^, whereas the majority of neoplasms in birds and reptiles were lymphosarcomas^25^. Dogs exposed to passive smoking and air pollution may develop cancer^114, 115^. Numerous other carcinogens in the environment and diet are known to affect malignancy and neoplasia prevalence in several species^38, 39, 116–118^, so it would be useful to know the impact of these carcinogens on malignancy and neoplasia across species. More environmental toxins may be found in the tissues of animals living in lower productivity habitats and/or in higher levels of the food pyramid^32, 39, 119^, which could explain the observed higher malignancy and neoplasia prevalence in lower productivity habitats (Fig. 2 & 3) as well as the higher cancer prevalence across tissues in higher trophic levels (Fig. 4 & 5; Supp. Fig. 1; Table 2).

We expected what type of food different species would predominantly consume rather than what they occasionally would consume to affect their evolution and development the most. Therefore, we classified species in this database according to their primary diet. Malignancy and neoplasia prevalence, however, may also depend on whether species are monotrophs or polytrophs (e.g., generalists). Therefore, malignancy and neoplasia prevalence could be compared with the diversity of food that an organism eats, although we do not have data on the whole range of foods that every species eats or a clear biological hypothesis about which group, monotrophs or polytrophs, may have higher cancer prevalence.

Future studies can test whether the correlations between higher trophic levels and cancer prevalence (Fig. 4 & 5; Supp. Fig. 1; Table 2) are also present when comparing carnivorous physiological traits and cancer prevalence. In humans, the risk of developing breast and colorectal cancer is higher in people that mainly work at night^49, 120^. Nocturnal and crepuscular species may have higher cancer prevalence than diurnal species, and nocturnal and crepuscular living is more typical of carnivores^121–123^. It may be that this association between carnivory and nocturnal or crepuscular living is partially responsible for the relationship we found between higher trophic levels and higher cancer prevalence (Fig. 4 & 5; Supp. Fig. 1; Table 2). The association between carnivores and cancer prevalence may also depend on the microbiome^124^, though little is currently known about the microbiome of most of the species in this dataset.

## Conclusion

To the extent that cancer has been an important selective factor on the survival and reproduction of a species, it would be expected that natural selection blunts any associations between ecological factors and cancer prevalence. Tradeoffs in responding to different selective pressures in an organism’s ecology, however, may constrain the ability of a species to effectively suppress the development of neoplasia. This study found that malignancy and neoplasia prevalence across tissues was correlated with trophic levels across vertebrates, even when applying Bonferroni corrections for multiple testing (Table 2). Future studies should distinguish whether this pattern exists due to bioaccumulation of toxins in high trophic levels, due to the consumption of meat per se, or both. An additional hypothesis for potential testing would be the effect of insect-derived chemicals, such as chitin, on the low cancer incidence in invertivores. Species with high inferred metabolic rates and in high productivity habitats have lower neoplasia prevalence but these relationships are not significant after Bonferroni corrections, and the reasons for those potential relationships remain unknown. These results suggest that species with higher metabolic rates and in higher productivity habitats might have more somatic resources with which to invest in cancer defences. A complete theory of metabolic ecology would help distinguish between these possible mechanisms and this knowledge could possibly be used to decrease the burden of cancer for animals managed in human care, free ranging animals, and humans.

## Supporting information

Supplementary_Extended References

Supplementary Data_P values

## Acknowledgements

Thanks to all of the pathologists, veterinarians, and staff at the zoos, aquariums, and veterinary hospitals for contributing to the data collection by diagnosing malignancy and neoplasia prevalence. Specifically, we would like to acknowledge the following institutions: Akron Zoo, Atlanta Zoo, Audubon Nature Institute, Bergen County Zoo, Birmingham Zoo, Buffalo Zoo, Capron Park Zoo, Central Florida Zoo, Dallas Zoo, El Paso Zoo, Elmwood Park Zoo, Fort Worth Zoo, Gladys Porter Zoo, Greensboro Science Center, Henry Doorly Zoo, Utah’s Hogle Zoo, Jacksonville Zoo, John Ball Zoo, Los Angeles Zoo, Louisville Zoo, Mesker Park Zoo, Miami Zoo, Oakland Zoo, Oklahoma City Zoo, Philadelphia Zoo, Phoenix Zoo, Pueblo Zoo, San Antonio Zoo, Santa Ana Zoo, Santa Barbara Zoo, Sedgwick County Zoo, Seneca Park Zoo, The Brevard Zoo, The Detroit Zoo, The Oregon Zoo, and Toledo Zoo. Thanks to Diego Mallo, Walker Mellon, and Rebecca Belshe for help with the statistical analyses. Thanks to Valerie Harris for contributing to the collection of life history data. Thanks to Andrew Beckerman and Michael Angilletta for helpful discussions on ecological relationships between species, food webs, and the metabolic theory of ecology. This work was supported in part by NIH grants U54 CA217376, U2C CA233254, P01 CA91955, and R01 CA140657 as well as CDMRP Breast Cancer Research Program Award BC132057 and the Arizona Biomedical Research Commission grant ADHS18-198847. The findings, opinions and recommendations expressed here are those of the authors and not necessarily those of the universities where the research was performed or the National Institutes of Health.

## Author Contributions

S.E.K. conceived and designed the study comparing cancer data with trophic levels, habitats, metabolic rates, and species body temperatures in February 2020. S.E.K. collected data on trophic levels, habitats, number of habitats, habitat productivity, average body temperatures, inferred metabolic rates, analysed the data, and wrote the first draft of the manuscript. T.M.H. helped in identifying the diet of animals in zoos. M.M.G., E.G.D., and T.M.H. helped in the collection and coordination of cancer data across institutions. S.M.R., Z.C., and A.M.B., provided data on malignancy and neoplasia prevalence, necropsies for each species, and species adult weight. A.A. and C.C.M. provided research supervision, helpful discussions, comments, and guidance throughout the project. All authors edited and approved the final version of the manuscript.

## Supplementary Figures

**Supplementary Figure 1. Neoplasia prevalence in relation to trophic levels in mammals versus non-mammals.** Neoplasia prevalence is always higher in higher trophic levels except for gastrointestinal neoplasia prevalence in mammals (Panel C). Each dot is a species, and each trophic level consists of ≥10 species. The number (N) of species in each trophic level is indicated above each trophic level. The horizontal black line in each category shows the median neoplasia prevalence in that category. We provide *P*-values between categories that are significantly different, and for all comparisons in the supplementary data and Table 2.

**Supplementary Figure 2. Neoplasias (B) but not malignancies (A), are more prevalent in species living in three or more habitats.** The y axes show malignancy prevalence across tissues (A), neoplasia prevalence across tissues (B), gastrointestinal malignancy prevalence (C), gastrointestinal neoplasia prevalence (D). Each dot is a species, and each category on the x axis consists of ≥10 species. The number of species (N) in a category is noted above each category. The horizontal black line in each category shows the median malignancy or neoplasia prevalence in that category. The x axis is classified as a categorical variable. We provide *P*-values between categories that are significantly different, and for all comparisons in the supplementary data and Table 2.

**Supplementary Figure 3. Malignancy prevalence across tissues is not significantly correlated with inferred metabolic rates.** The formula M ^−1/4^ • e ^−E/kT^ multiplied by bo is a measure of metabolic rate for each species. The bo value [bo = a coefficient independent of body size and temperature from Savage et al. 2004^58^] is unknown for each species, so we measured the M ^−1/4^ • e ^−E/kT^ values for each species (where M = mass, E = 0.65 eV, k = 8.62 • 10^−5^ eVK^−1^, and T = temperature in Kelvin)^56^. Each dot is a species.

**Supplementary Figure 4. Malignancy and neoplasia prevalence are not significantly associated with average body temperature (in Kelvin) across the tree of life.** The y axes show malignancy prevalence across tissues (A: F-statistic: 0.80 on 1 and 78 DF, Lambda: 0.80) and neoplasia prevalence across tissues (B: F-statistic: 0.24 on 1 and 78 DF, Lambda: 0.85). Each dot is a species.

**Supplementary Figure 5. M** ^−1/4^ **• e ^−E/kT^ (which is analogous to metabolic rate) to the power of 0.1 is lower in higher trophic levels.** Each dot is a species. This measure (M^−1/4^ • e ^−E/kT^) multiplied by bo is a measure of metabolic rate for each species. We do not know the bo value [bo = a coefficient independent of body size and temperature from Savage et al. 2004^58^] for each species, so we simply measured the M ^−1/4^ • e ^−E/kT^ values for each species (M = mass, E = 0.65 eV, k = 8.62 • 10^−5^ eVK^−1^, and T is temperature in Kelvin)^56^. The number (N) of species in each trophic level is indicated above each trophic level. The horizontal black line in each category shows the median. F-statistic: 1.84 on 2 and 67 DF, Lambda: 0.77, R²: 0.05. We only provide *P*-values between categories in the figures if the comparisons are significantly different. All *P*-values for the category comparisons are available in the supplementary data.

**Supplementary Figure 6. The majority of carnivores in our database live in many different habitats, whereas the majority of herbivores live in one or two habitats.** N shows the number of species in each level. The percentage shows the number of species in a specific habitat range out of the total species in that trophic level • 100%. Percentages in bold show the two highest percentages of species in each trophic level.

**Supplementary Figure 7. The majority of herbivores and invertivores live in high productivity habitats, whereas the majority of primary and secondary carnivores live in low productivity habitats.** Each dot is a species. We used minimal jitter to better visualise individual dots. Numbers show the number of species in each level. Numbers are shown only in the three columns, e.g. habitats where we have ≥10 species in our database. The percentage shows the number of species in a specific habitat out of the total species in that trophic level • 100%. Percentages in bold show the two highest values in each habitat per trophic level.

