## Supplementary figures and images for "The ecology of cancer prevalence across species: Cancer prevalence is highest in desert species and high trophic levels"

### Supplementary Figure 1

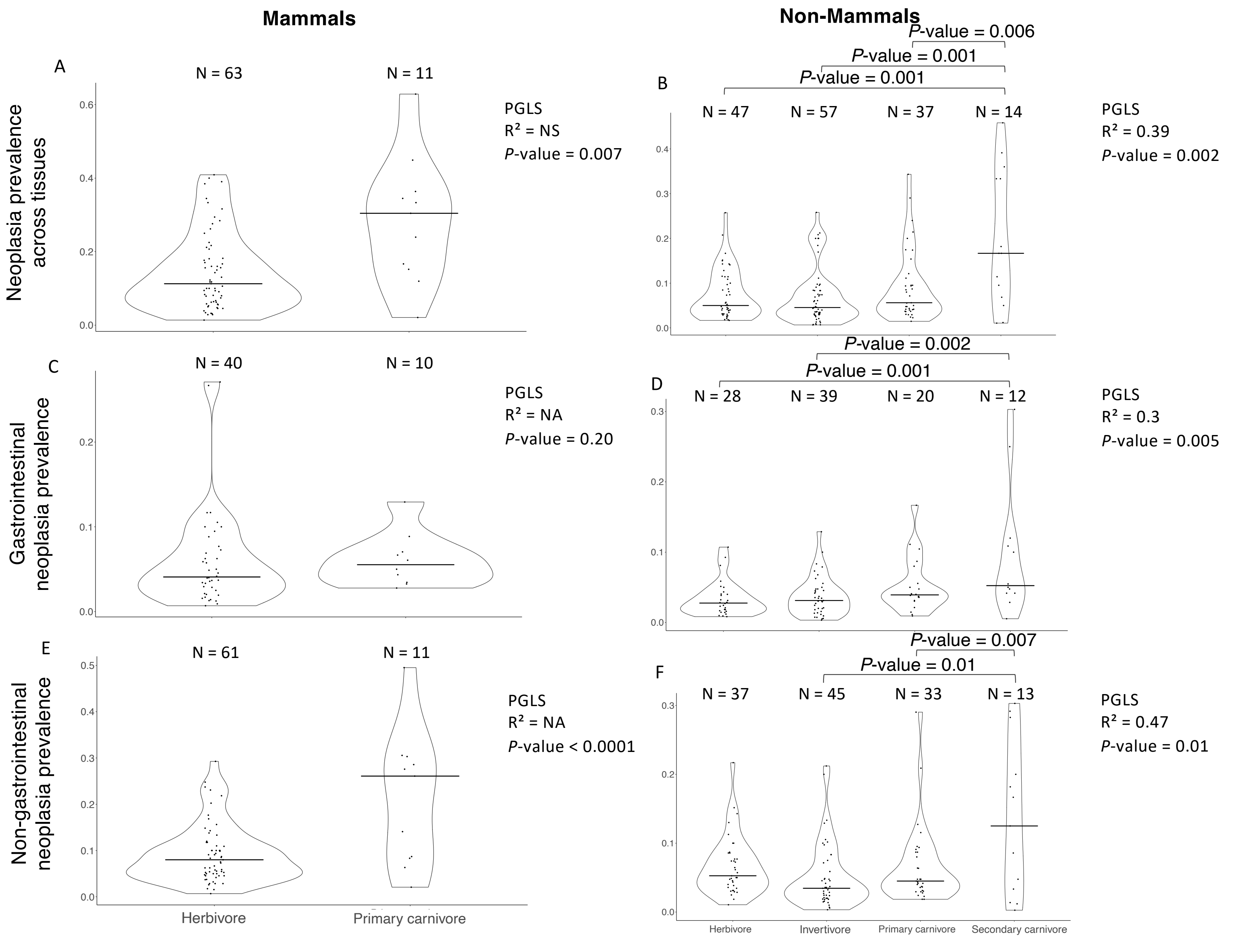

### Supplementary Figure 2

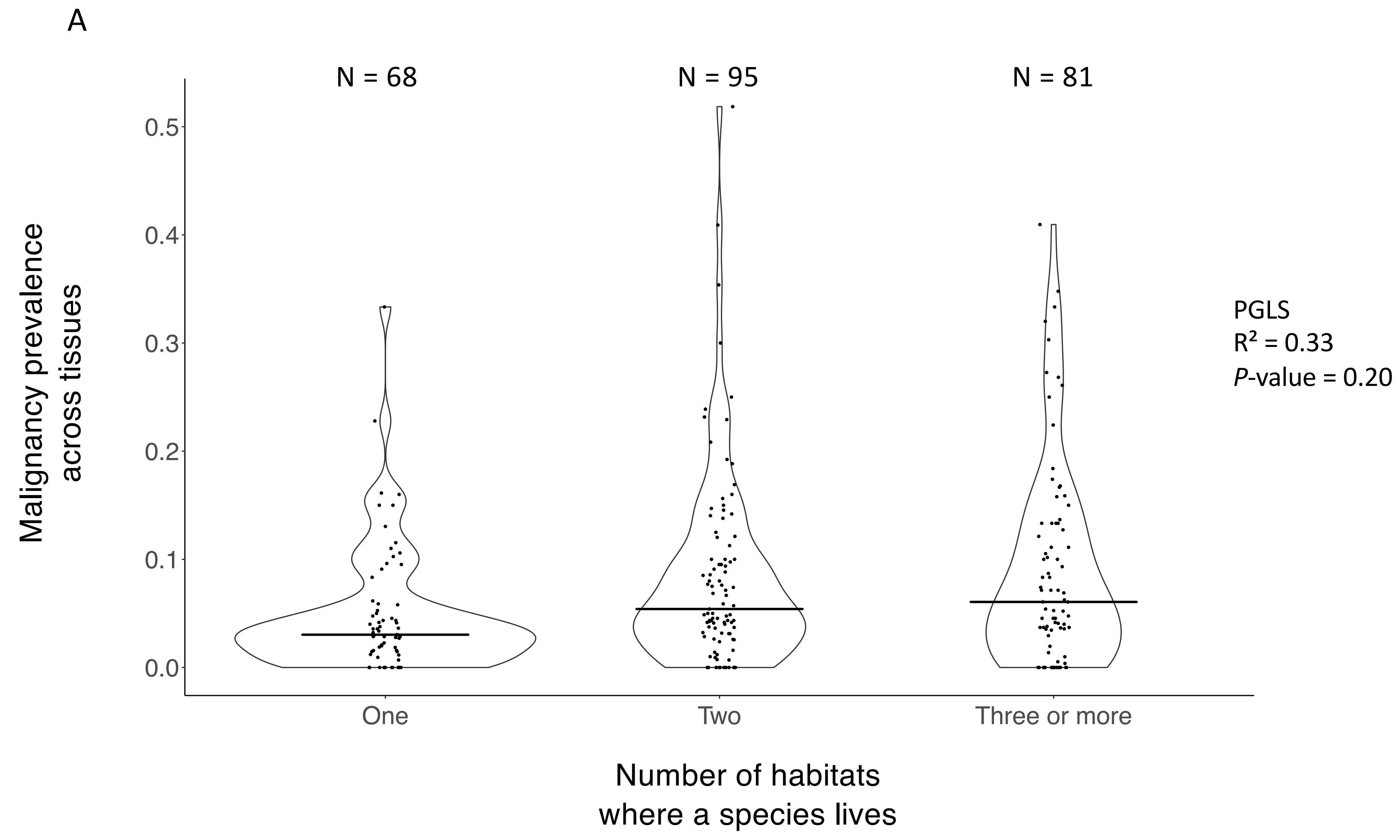

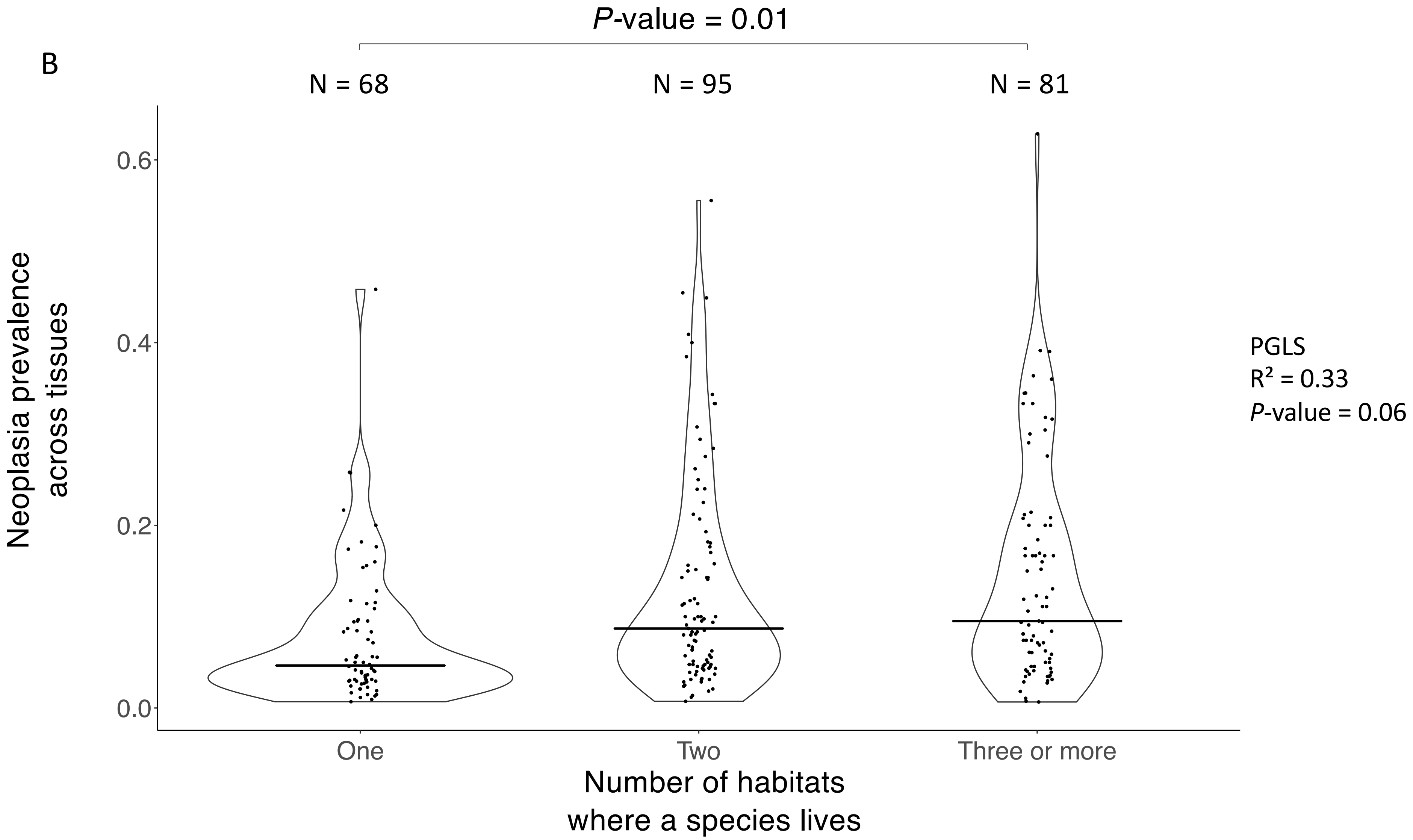

### Supplementary Figure 3

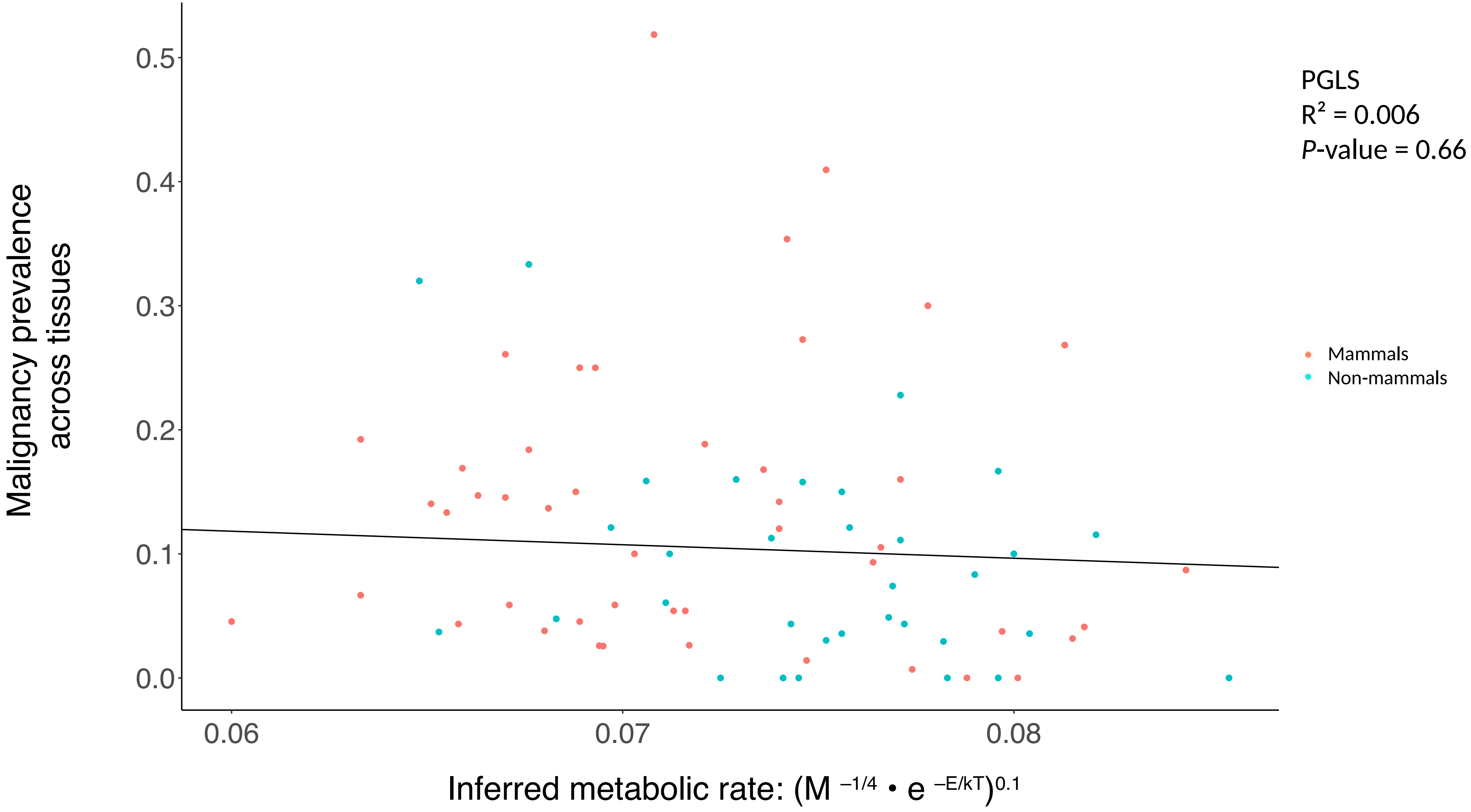

### Supplementary Figure 4

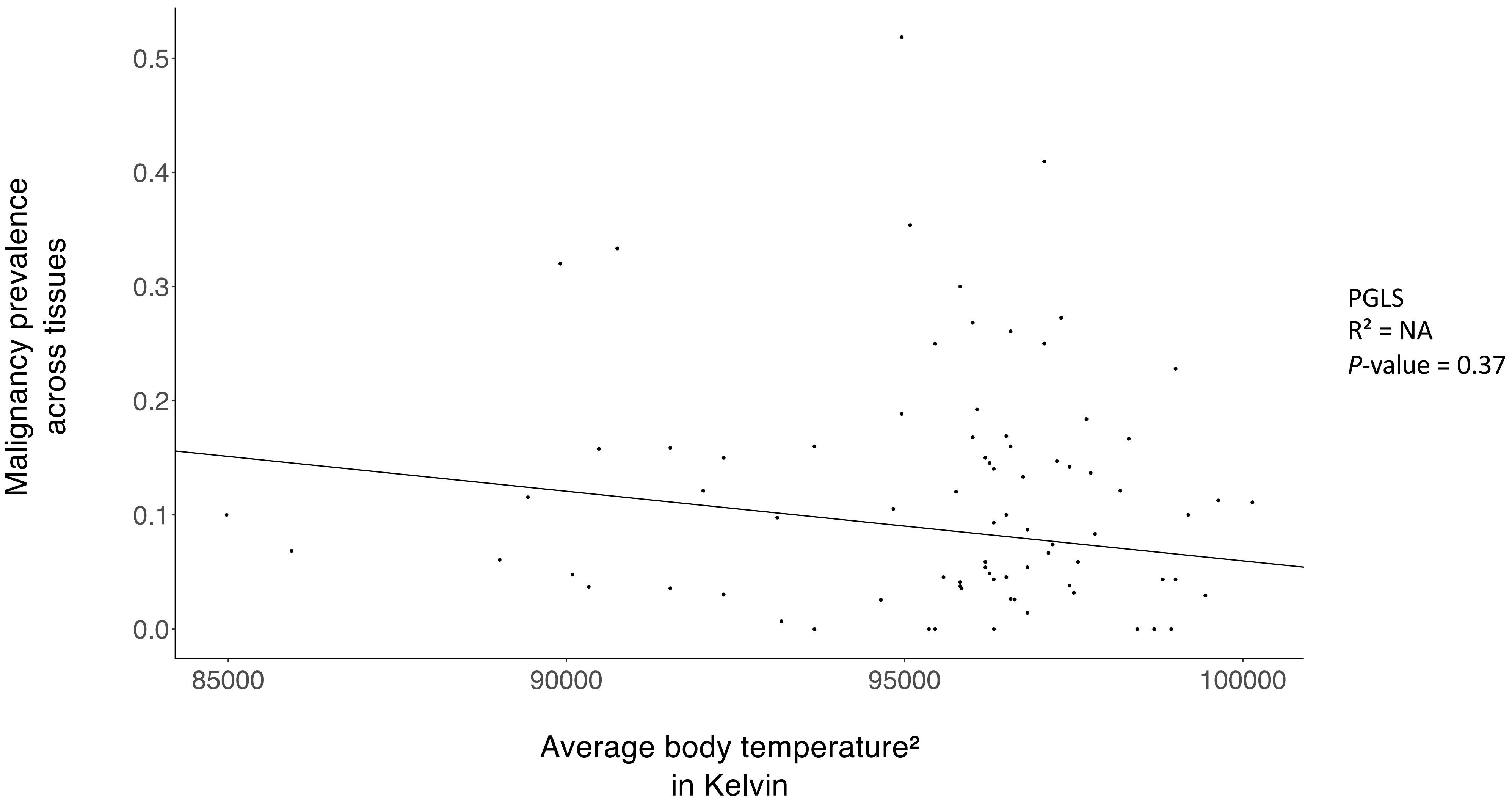

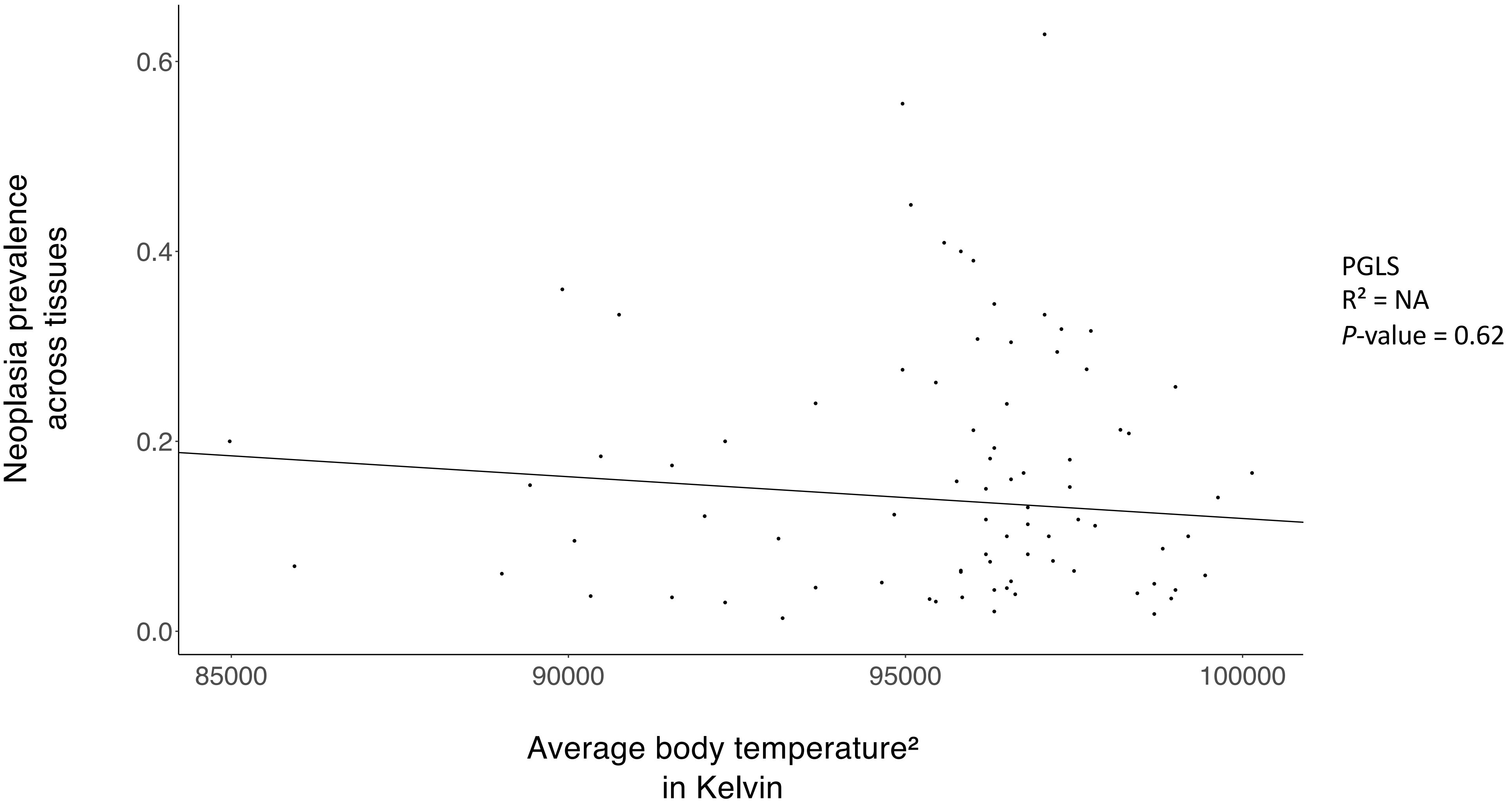

### Supplementary Figure 5

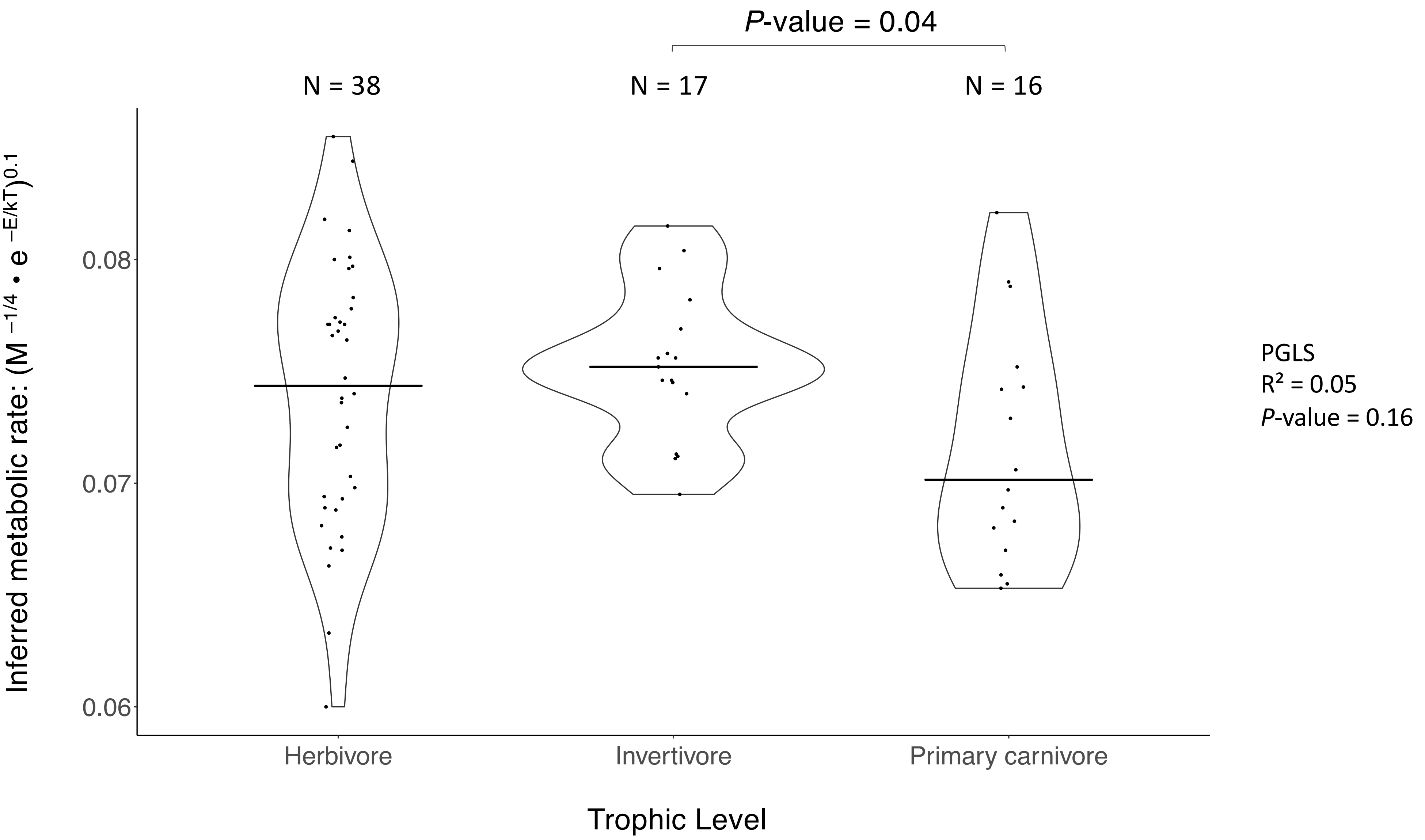

### Supplementary Figure 6

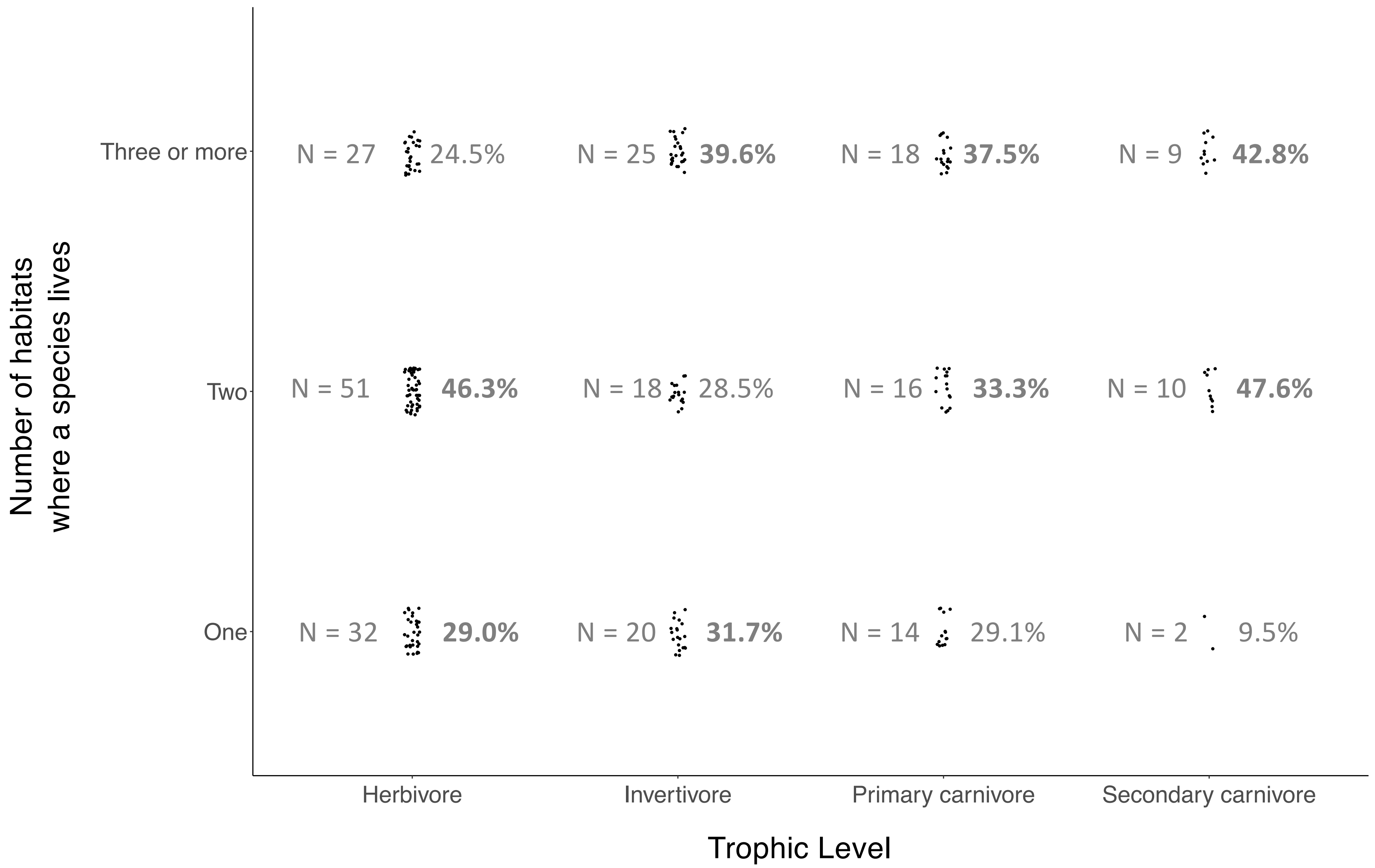

### Supplementary Figure 7

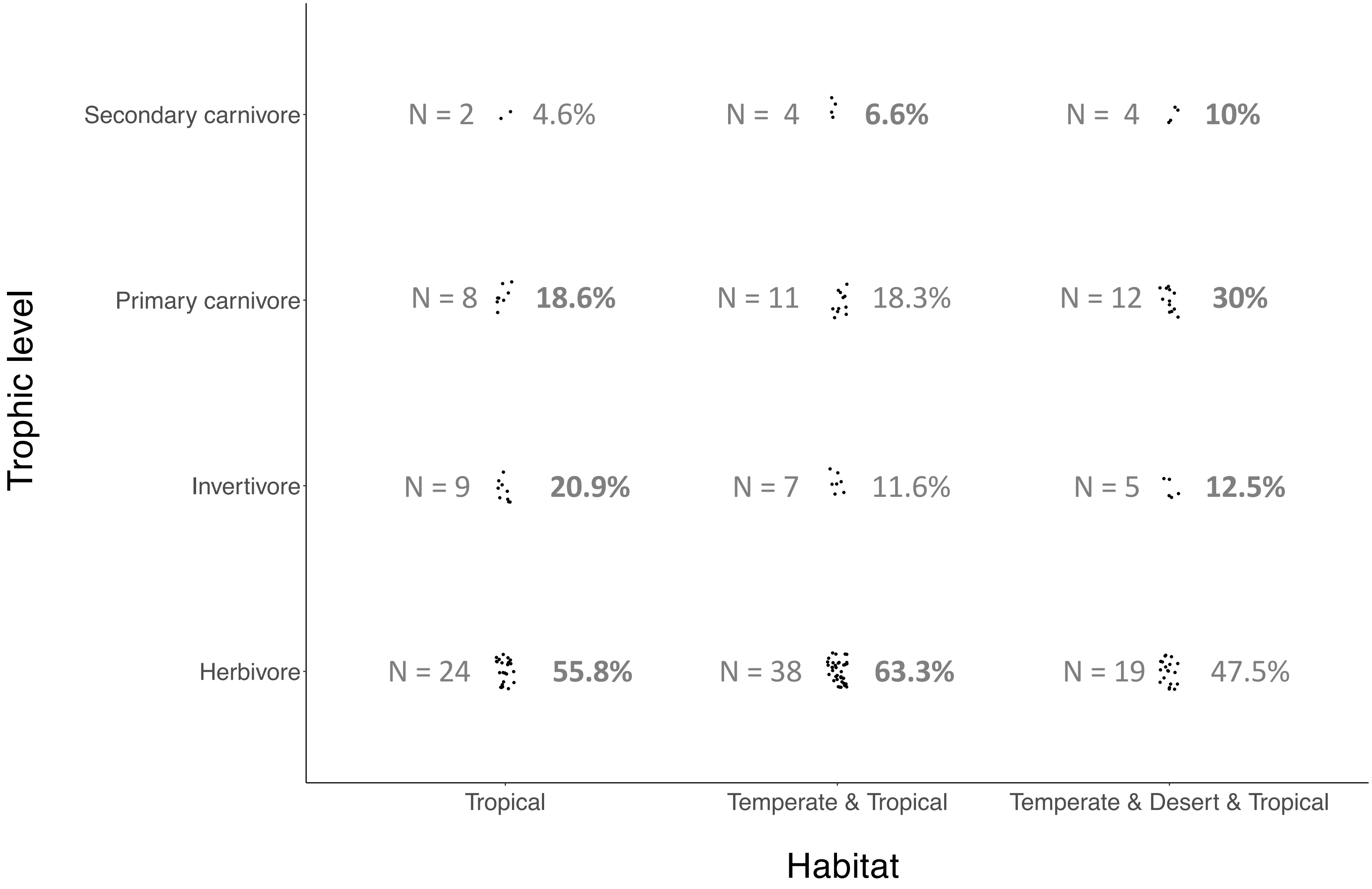
